# Fine mapping spatiotemporal mechanisms of genetic variants underlying cardiac traits and disease

**DOI:** 10.1101/2021.09.01.458619

**Authors:** Matteo D’Antonio, Timothy D. Arthur, Jennifer P. Nguyen, Hiroko Matsui, Agnieszka D’Antonio-Chronowska, Kelly A. Frazer

**Affiliations:** Department of Pediatrics, University of California San Diego, La Jolla, CA, 92093, USA; Department of Biomedical Informatics, University of California, San Diego, La Jolla, CA, 92093, USA; Biomedical Sciences Graduate Program, University of California, San Diego, La Jolla, CA 92093, USA; Bioinformatics and Systems Biology Graduate Program, University of California, San Diego, La Jolla, CA, 92093, USA; Institute of Genomic Medicine, University of California San Diego, 9500 Gilman Dr, La Jolla, CA, 92093, USA

## Abstract

The causal variants and genes underlying thousands of cardiac GWAS signals have yet to be identified. Here, we leveraged spatiotemporal information on 966 RNA-seq cardiac samples and performed an expression quantitative trait locus (eQTL) analysis detecting eQTLs considering both eGenes and eIsoforms. We identified 2,578 eQTLs associated with a specific developmental stage-, tissue- and/or cell type. Colocalization between eQTL and GWAS signals of five cardiac traits identified variants with high posterior probabilities for being causal in 210 GWAS loci. Pulse pressure GWAS loci were enriched for colocalization with fetal- and smooth muscle- eQTLs; pulse rate with adult- and cardiac muscle- eQTLs; and atrial fibrillation with cardiac muscle- eQTLs. Fine mapping identified 79 credible sets with five or fewer SNPs, of which 15 were associated with spatiotemporal eQTLs. Our study shows that many cardiac GWAS variants impact traits and disease in a developmental stage-, tissue- and/or cell type-specific fashion.

## Introduction

Genome-wide association studies (GWAS) have identified thousands of loci associated with cardiac traits and diseases ^1^, but for the vast majority of these associations the underlying causal variants and genes have not yet been delineated. Many approaches have been developed to identify causal variants and genes, including annotating GWAS loci with their closest gene ^1^, prioritizing variants that overlap regulatory elements active in cardiac tissues ^2,3^, and integrating GWAS variants with adult cardiac eQTL (expression quantitative trait loci) signals ^4^. However, these approaches have been limited in part due to the fact that regulatory elements can regulate the expression of multiple genes over hundreds of kilobases ^5,6^; and regulatory variants frequently work in a spatiotemporal context not captured in previous eQTL analyses ^7–11^. For some GWAS traits and diseases efforts have been made to fine map each GWAS locus and identify potential associations between genetic variation, gene expression and disease^12–14^; however, the cardiac field still lacks a resource for conducting in depth annotations of GWAS loci. Here, we integrate GWAS signals for multiple cardiac traits with gene expression data for different cardiac developmental stages, tissues, and cell types across hundreds of individuals and demonstrate the power of this comprehensive resource for fine mapping causal regulatory variants and understanding the molecular mechanisms underlying cardiovascular GWAS traits and disease.

Gene expression has long been known to be regulated in a spatial (organ, tissue or cell type) and temporal (fetal-like and adult) specific manner ^15–22^, indicating that cardiac regulatory variants function, and hence affect the expression of a gene and its associated cardiac traits and disease, across a range of developmental stages and in different cellular contexts ^15–17,23–26^. With the development of cell type deconvolution techniques ^27,28^, bulk RNA-seq enables the characterization of cell type-specific gene expression as well as the expression of both genes and associated isoforms ^24,29,30^. Furthermore, the GTEx consortium has generated bulk RNA-seq from hundreds of adult cardiac samples from multiple tissue types and whole genome-sequenced the donors ^25^. Although several fetal-associated factors, such as low birth weight, maternal preeclampsia, under- and malnutrition and oxidative stress in utero, have been associated with increased risk of developing cardiovascular disease as adults ^31,32^, large numbers of human fetal cardiac specimens are not readily available, and functional genetic variation in fetal heart cell types remains largely unstudied. To overcome ethical and availability issues associated with the use of fetal samples, iPSCORE has developed strategies to employ induced pluripotent stem cell (iPSC) derived cardiovascular precursor cells (iPSC-CVPCs) from hundreds of whole-genome sequenced individuals to study early developmental cardiac traits and disease ^29,33–35^. We and others have shown that iPSC-CVPCs display epigenomic and transcriptomic properties similar to that of fetal cardiac cells ^29,36–41^, and their differentiation results in both cardiomyocytes and epicardium-derived cells ^33^. We have also previously deconvoluted the proportions of cell types in ~ 1,000 bulk RNA-seq samples from fetal-like iPSC-CVPCs ^33^, adult healthy heart and arteria ^42^ and heart failure ^43^, and demonstrated the power of this approach for examining the reactivation of fetal-specific genes and isoforms during heart failure ^29^.

In this study, we show that the combined deconvoluted fetal-like iPSCORE and adult GTEx ^25,33^ cardiac expression datasets enables the genome-wide mapping of regulatory variant functions in a spatial-temporal specific manner. We performed an expression quantitative trait locus (eQTL) analysis on 966 deconvoluted bulk RNA-seq samples, including adult (atrium, ventricle, aorta and coronary artery) and fetal-like (iPSC-CVPCs) tissues, and identified cardiac eQTL signals associated with 11,692 eGenes and 7,165 eIsoforms. Less than half of the eIsoforms shared the same eQTL signal with their associated gene, indicating that molecular mechanisms underlying the associations of genes and their isoforms with cardiac traits and disease are different in many cases. By leveraging information about the source and cellular composition of each sample, we were able to detect more than 2,578 eQTLs that function in a spatiotemporal manner. We exploited the cardiac eQTLs to investigate the molecular underpinnings of five cardiac traits and diseases from the UK BioBank and found that many of the cardiac GWAS signals colocalized with eQTL signals ^44^ that function in a specific cardiac stage, organ, tissue and/or cell type context. We also observed that three of the cardiac traits were enriched for eQTLs that function in specific spatiotemporal contexts, including pulse pressure with fetal-like-, arteria- and smooth muscle cell-eQTLs; pulse rate with adult-, heart- and atrial-eQTLs; and atrial fibrillation with left ventricle- and cardiac muscle cell-eQTLs. We used the colocalized eQTL and GWAS signals for fine mapping, which allowed us to identify a potential causal variant for 210 cardiac GWAS loci. Overall, our study serves as a comprehensive resource mapping cardiac regulatory variants that function in spatiotemporal context-specific manners to alter gene expression and affect cardiac traits, thereby providing potential molecular mechanisms underlying the associations of hundreds of GWAS loci with cardiac traits and disease.

## Results

To investigate associations between genetic variation and cardiac gene expression we obtained RNA-seq for 180 fetal-like iPSC-CVPCs (derived from 149 iPSC lines) from 139 individuals included in the iPSCORE collection ^33^ and integrated with RNA-seq data for 786 adult cardiac tissues, including atrial appendage, left ventricle, aorta and coronary artery from 352 individuals included in the GTEx Consortium ^42^ (Table S1). To map the regulatory effects of genetic variants on these fetal-like and adult cardiac tissues, we performed an eQTL analysis on all 966 samples using a linear mixed model (LMM) with developmental stage (iPSC-CVPC or adult), organ (arteria or heart), tissue (atrium, ventricle, aorta and coronary artery) and deconvoluted cell type proportions ^29^ as interaction terms. Among 19,586 expressed autosomal genes (TPM ≥ 1 in at least 10% samples), we identified at least one eQTL for 11,692 genes (eGenes, 59.7% of tested genes, Figure 1A, Table S2). By regressing out the genotype of each lead variant, we observed that, on average, each eGene had 1.54 eQTLs (range: 1-6), in line with what has recently been reported by GTEx ^42^. Specifically, we obtained conditional eQTLs for 4,394 eGenes (37.6% of all eGenes), including 1,315 with two conditional eQTLs, 395 with three, 160 with four and 74 with five (Figure 1A). We also examined 37,032 autosomal isoforms (corresponding to 10,337 genes; at least two isoforms/gene with usage ≥ 10% in at least 10% samples) and identified 7,165 with at least one eQTL (eIsoforms, 19.3% of all tested isoforms, corresponding to 3,847 genes), including 988 with one or more conditional eQTL (Figure 1A). We identified fewer eIsoforms than eGenes likely because of decreased power in detecting eQTL associations, caused by a more stringent multiple testing correction, as we tested twice as many isoforms than genes.

**Figure 1:**
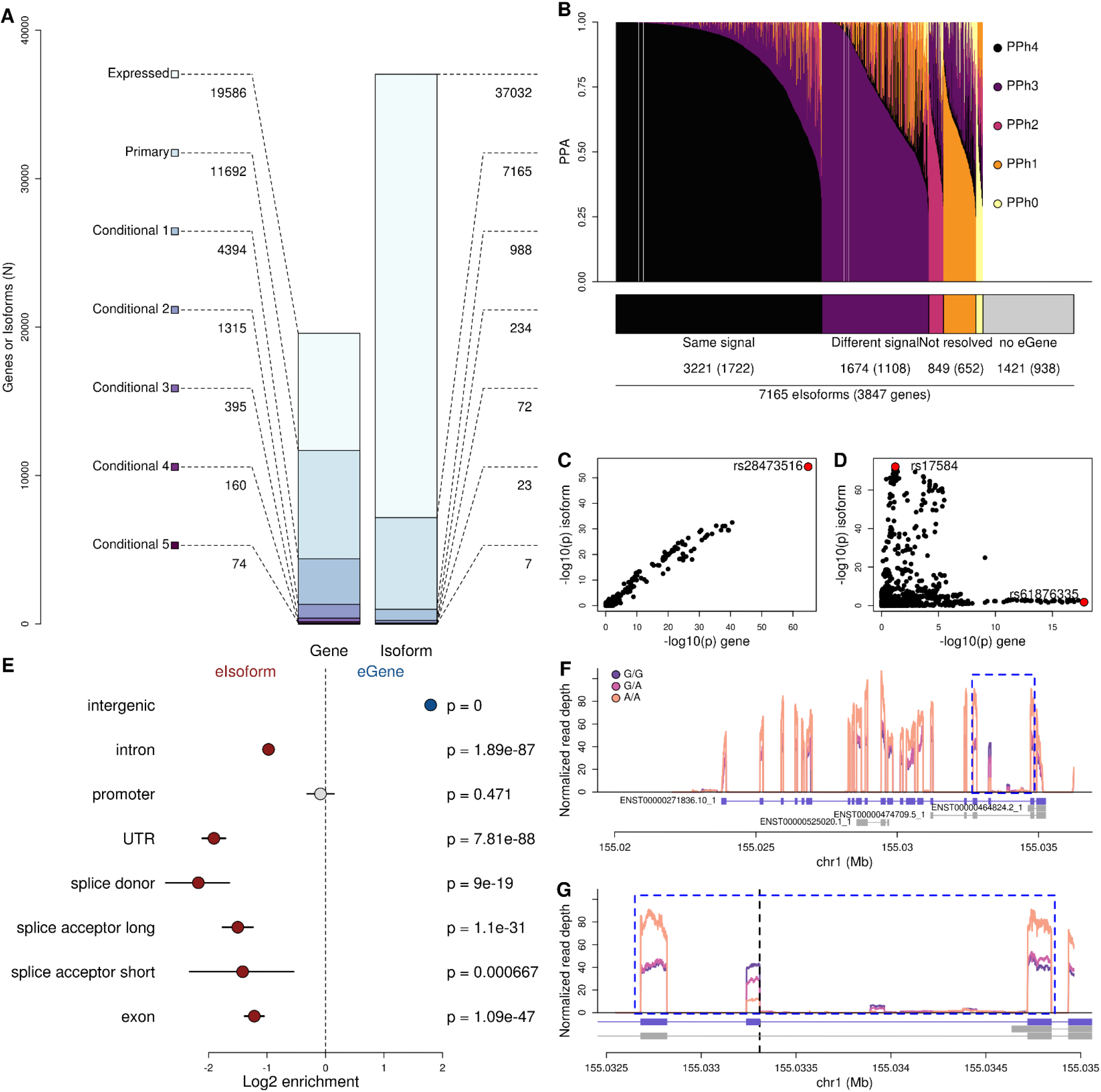
Gene and isoform eQTLs. (A) Barplot showing the number of eQTLs for eGenes (left) and eIsoforms (right). Colors represent whether each eGene or eIsoform does not have eQTLs (white), the number of eGenes with primary and conditional eQTLs (up to five conditional signals). (B) Barplot showing the distribution of PPA for each of the five colocalization hypotheses, as described by Giambartolomei et al. {Giambartolomei, 2014 #28}: 1) H0: neither trait (eQTL signal for eIsoform and its associated eGene) has a significant association at the tested locus; 2) H1: only the first trait (eIsoform) is associated; 3) H2: only the second trait (eGene) is associated; 4) H3: both traits are associated but the underlying variants are different; and 5) H4: both traits are associated and share the same underlying variants. All eIsoforms having H4 as the most likely hypothesis were considered as sharing the same eQTL signal with their associated eGene, whereas all the eIsoforms having H3 as the most likely hypothesis were considered as having a different eQTL signal than their associated eGene. All the colocalizations associated with hypothesis 0, 1 and 2 were likely underpowered, thus they were labeled as “not resolved”. The 1,421 eIsoforms whose associated gene did not have an eQTL were labeled as “no eGene”. (C,D) Examples of (C) eQTL signal for an eGene (*B4GALT7*) that colocalizes with PPA = 1 with the eQTL signal of one associated eIsoform and (D) eQTL signal for an eGene (*RNH1*) that does not colocalize with the eQTL signal of one associated eIsoform. In each plot, X axis represents the −log_10_ (p-value) for the associations between the genotype of each tested variant and gene expression, whereas the Y axis shows the −log_10_ (p-value) for the associations between the genotype of each tested variant and isoform use. In red the lead variants are shown (two in panel C, as eGene and eIsoform have different associations, one in panel B, as the eGene and eIsoform signal colocalize). (E) Enrichment of eGenes compared with eIsoforms for overlapping intergenic regions, introns, promoters, UTRs, splice donor sites, splice acceptor sites (short = the first 5 nucleotides upstream of the splice site; long = the first 100 bp) and exons. P-values were calculated using Fisher’s exact test. Points (blue = enriched for eGenes; red = enriched for eIsoforms; gray = not significant) represent log_2_ enrichment and horizontal lines represent 95% confidence intervals calculated using the *fisher.test* function in R. (F,G) Median normalized read depth signal of *ADAM15* gene expression levels in iPSC-CVPCs. Different colors represent the genotypes of the lead eVariant for isoform ENST00000271836.10_1 (rs11589479, G>A). The blue rectangle in (F) is enlarged in (G). rs11589479 overlaps the splice donor site for exon 19 and its position is shown as a vertical dashed line in (G). The plots show that the exon whose splice site is affected by rs11589479 becomes expressed at lower levels when the variant is heterozygous or homozygous alternative, as it disrupts the splice site.

### Different mechanisms underlie eQTLs for genes and isoforms

We investigated the extent to which underlying genetic mechanisms differ between eQTL signals for gene expression and isoform usage. Of the 3,847 genes associated with the 7,165 eIsoforms, 2,909 were themselves eGenes (corresponding to 5,744 eIsoforms), while 938 were not eGenes (corresponding to 1,421 eIsoforms). For the 2,909 eGenes with eIsoforms, to examine if the same genetic variants were associated with both gene expression and isoform use, we performed a colocalization analysis ^44^ (a Bayesian approach that provides posterior probabilities of signals for two traits at one locus (trait 1 = eIsoform; trait 2 = eGene) for five hypotheses: H0) neither trait is associated; H1) only trait 1 is associated; H2) only trait 2 is associated; H3) both traits are associated, but with different underlying causal variants; and H4) both traits are associated with the same underlying causal variants). For 3,221 eIsoforms (56.2% of the 5,744 associated with eGenes) the most likely hypothesis was 4, indicating that the eIsoform and eGene eQTL signals share the same causal variant (Figure 1B, Table S3). For example, *B4GALT7* and its isoform (ENST00000029410.10_2) shared a common eQTL signal, with their lead variant (rs28473516) having >99% PPA of being causal for both gene expression and isoform use (Figure 1C). On the other hand, 1,674 (29.1%) of the eIsoforms had a different signal than their associated eGene. For example, the lead variant for *RNH1* (rs61876335) was located ~6 kb downstream of the 3’ end of the gene, whereas the lead variant for its isoform (rs17584, ENST00000397604.7_2), which is not in LD with rs61876335 (r^2^ = 0.019 in EUR individuals), was a synonymous coding variant (Figure 1D). For the remaining 849 (14.7%) eIsoforms, the eQTL signals for the eIsoform (hypothesis 1), for the eGene (hypothesis 2) or for both (hypothesis 0) were likely underpowered. These results show that at least 29.1% of eIsoforms had a different signal than their associated eGene, suggesting that different mechanisms underlie the association of genetic variation with gene expression and isoform use.

To test if eQTLs for genes and isoforms have different properties, we investigated their overlap with gene bodies and promoters. We observed that gene eQTLs are more likely to occur in intergenic regions than isoform eQTLs (p ≈ 0), whereas isoform eQTLs are more likely to overlap gene bodies, including introns (p = 1.9 × 10^-87^), UTRs (p = 7.8 × 10^-88^), splice donor sites (p = 9.0 × 10^-19^), splice acceptor sites (p = 1.1 × 10^-31^) and exons (p = 1.1 × 10^-47^, Figure 1E), indicating that isoform eQTLs are more likely to influence transcript stability than regulatory elements. For example, we found that the lead eVariant (rs11589479) for an *ADAM15* isoform (ENST00000271836.10_1: the most common isoform of *ADAM15*) is a G>A substitution that likely disrupts the splice site for exon 19 (Figure 1F,G). *ADAM15* encodes a disintegrin and metalloprotease involved in cell-cell and cell-matrix interactions and has an established role in inflammation and angiogenesis ^45^. Of note, rs11589479 did not colocalize with the primary eQTL signal for *ADAM15* expression. We observed that *ADAM15* exon 19 was expressed at lower levels in samples carrying heterozygous or homozygous alternative alleles for rs11589479 and that the overall expression of exon 19 was reduced by ~80% in homozygous alternative samples compared with the surrounding exons, whereas in samples carrying homozygous reference alleles for rs11589479 the expression of exon 19 was comparable with the surrounding exons. These data show that the molecular mechanisms underlying the associations of genes and their isoforms with cardiac traits and disease will be different in many cases, with gene expression being likely associated with regulatory variants at promoters and enhancers and isoform usage associated with variants that affect post-transcriptional modifications.

### Mapping spatiotemporal cardiovascular eQTLs

To identify eQTLs that function in a spatiotemporal manner, we compared the eQTL signal between samples at each stage (iPSC-CVPC or adult) or annotated as a specific organ (arteria and heart) or tissue (atrium, ventricle, aorta and coronary artery) against all the other samples using an interaction test between genotype and each of these features ^46^. From this analysis, we classified eQTL signals as shared, specific or associated (Figure 2A-E, Figure S1, Figure S2, Table S4). For example, for *RPS26* we did not observe a significant interaction between the genotype of the lead eQTL signal variant (rs1131017) and stage (β = 0.002, p = 0.21, Figure 2C), therefore we annotated this eQTL as “shared” between both stages. Conversely, the eQTL signals for *DDTL* and for *ADAM15* transcript ENST00000271836.10_1 showed a significant interaction between genotype (rs9612520 and rs11589479, respectively) and stage (β = −0.90, p = 2.0 × 10^-42^ for *DDTL;* β = −1.09, p = 2.6 × 10^-7^ for *ADAM15*, Figure 2D,E), suggesting that their expression is differentially associated with genotype between iPSC-CVPC and adult heart. However, these two eQTL signals show substantial differences. The genotype of rs9612520 is not associated with *DDTL* in adult cardiac samples (β = 0, p = 0.90), suggesting that this eQTL signal is iPSC-CVPC-specific, whereas the eQTL signal for *ADAM15* is significant in both iPSC-CVPC (β = −2.32, p = 2.7 × 10^-22^) and adult heart (β = −0.92, p = 8.0 × 10^-14^), when tested independently. Since the signal in iPSC-CVPC is stronger than adult, we labeled this eQTL as “iPSC-CVPC-associated”. We found 814 stage-eQTL signals (combined -specific and -associated) for eGenes and 297 for eIsoforms (Bonferroni-corrected p < 0.05). Of these, the majority (620 eGenes and 191 eIsoforms) had adult-specific eQTL signals; 105 eGenes and 76 eIsoforms had associated eQTLs (i.e., in both stages but with significantly different effect sizes); and 89 eGenes and 30 eIsoforms had iPSC-CVPC-specific eQTL signals. We also observed organ- and tissue-specific or -associated eQTL signals for 1,246 eGenes and 350 for eIsoforms (Figure 2A,B, Figure S2). Since the same eQTL signal may be associated with both stage and tissue or organ, in total our interaction eQTL approach identified 1,665 eQTL signals for eGenes and 565 for eIsoforms associated with cardiac stage, organ and/or tissue.

**Figure 2:**
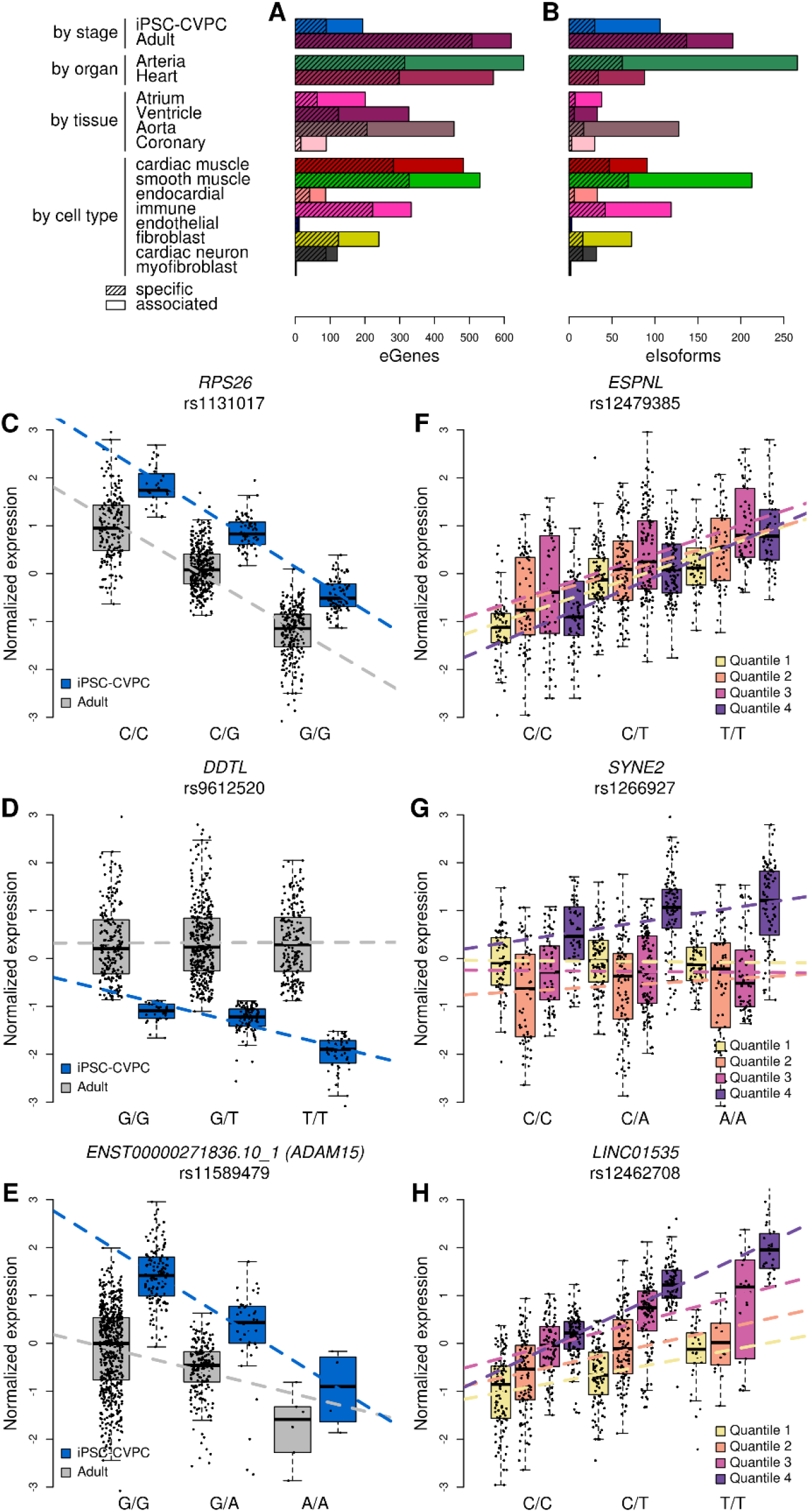
Stage, organ, tissue and cell type eQTLs. (A,B) Barplots showing the number of (A) eGenes and (B) eIsoforms associated with: cardiac stage (iPSC-CVPC or adult); organ (arteria or heart); tissue (atrial appendage, left ventricle, aorta or coronary artery); and cell type. Non-hatched bar sections represent eQTLs that are associated with the indicated stage, organ, tissue or cell type, and hatched sections represent specific eQTLs. (C-E) Examples of three association types between eQTLs and cardiac stage. For each eGene, boxplots describe the normalized expression in iPSC-CVPCs (blue) and all other samples (i.e., adult cardiac samples; gray), grouped by genotype. The panels show examples of: (C) an eGene whose eQTL is shared across both cardiac stages; (D) an iPSC-CVPC-specific eQTL: the association between genotype and gene expression is only present in iPSC-CVPCs; and (E) iPSC-CVPC-associated eQTL: while the genotype is associated with gene expression in both iPSC-CVPCs and the adult samples, the eQTL is significantly stronger in iPSC-CVPCs. All five possible association types are shown in Figure S1. (F-H) Examples of associations between eQTLs and cell types. For each eGene, boxplots describe the normalized expression divided into four quartiles according to their cardiac muscle proportion (yellow = low; purple = high), grouped by genotype. The panels show examples of: (F) an eGene whose eQTL is shared across cell types; (G) a cardiac muscle-specific eQTL: the association between genotype and gene expression is only present in the top quartiles; and (H) cardiac muscle-associated eQTL: while the genotype is associated with gene expression in all quartiles, the eQTL is significantly stronger in the top quartiles. All five possible association types are shown in Figure S3.

To assess the associations between eQTLs and cell types, we divided the 966 RNA-seq samples into quartiles according to their deconvoluted cell type populations, and for each cell type, tested the interaction between genotype and cell type proportions and compared the top and bottom quartiles (Figure 2F-H, Figure S3). We identified cases, such as *ESPNL*,where the interaction was not significant (β = −0.003, p = 1.0, Figure 2F), suggesting that these eQTLs were shared across cell types. For eQTL signals with a significant interaction, we compared the top and bottom quartile and annotated the eQTL as “cell type-specific” if only the expression levels in the samples included in the top quartile were associated with the genotype (for example, *SYNE2* and cardiac muscle, Figure 2G), and as “cell type-associated” if both the top and bottom quartile were significantly associated, but the signal in the top quartile was stronger (for example, *LINC01535*, Figure 2H) Using this method, we found 1,191 cell type-specific or -associated eQTL signals for eGenes and 364 for eIsoforms (Figure 2A,B). To validate this approach for identifying cell type-eQTLs (combined -specific and -associated), we tested the overlap between cardiac muscle-eQTLs with regulatory elements in nine cardiac cell types obtained from a single nuclei ATAC-seq (snATAC-seq) study on adult cardiac cells ^47^. We observed that cardiac muscle-eQTLs were more likely than expected to overlap regulatory elements enriched for being active in atrial and ventricular cardiomyocytes (p = 4.8 × 10^-14^ and p = 8.4 × 10^-9^, respectively, Figure S4, Table S5) and were less likely to overlap regulatory elements active in other cell types, including macrophages (p = 8.1 × 10^-16^), fibroblasts (p = 6.4 × 10^-10^) and adipocytes (p = 6.4 × 10^-3^). Overall, these data show that integrating stage, organ, tissue and cell type information with genotype and gene expression, we were able to determine the spatiotemporal context of 2,578 eQTL signals (Table S4).

### eQTL signals associated with multiple eGenes are enriched for being spatiotemporal regulated

Enhancers can regulate the expression of more than one gene ^6^, therefore we investigated how often eGenes in close proximity share the same eQTL signal. Using colocalization, we found that 2,778 eQTL signals were shared between two or more eGenes or eIsoforms from different genes (PPA > 0.8, range = 2-9 genes; mean = 2.21 ± 0.62, Table S6). We next investigated whether eQTL signals shared between multiple eGenes are enriched for being spatiotemporal regulated (i.e., associated with a cardiac stage-, organ-, tissue- or cell type). We found a significant positive association between the number of eGenes that share the same eQTL signal and the likelihood of their eQTL to be spatiotemporal regulated: adult (combined -specific and -associated) (p = 1.6 × 10^-17^, linear regression, Figure 3A), organ (arteria: p = 9.9 × 10^-8^; and heart: p = 5.5 × 10^-4^), tissue (ventricle: p = 0.023; and aorta: p = 4.8 × 10^-3^) and/or cell type (cardiac muscle: p = 1.1 × 10^-3^; smooth muscle: p = 9.5 × 10^-4^; immune cells: p = 3.1 × 10^-7^; and fibroblasts: p = 3.3 × 10^-12^). The difference in the enrichment between iPSC-CVPC and adult heart could suggest that fetal-eQTLs are less likely to be associated with multiple eGenes than adult-eQTLs. Furthermore, we observed that eGenes that share the same eQTL signals are significantly more likely than expected to be associated with the same stage, organ, tissue or cell type (Figure 3B). These results indicate that eVariants that are associated with multiple eGenes are enriched for being in regulatory elements that function in a temporal (in the adult, rather than in the fetal-like heart) and spatial (organ, tissue or cell type) specific manner.

**Figure 3:**
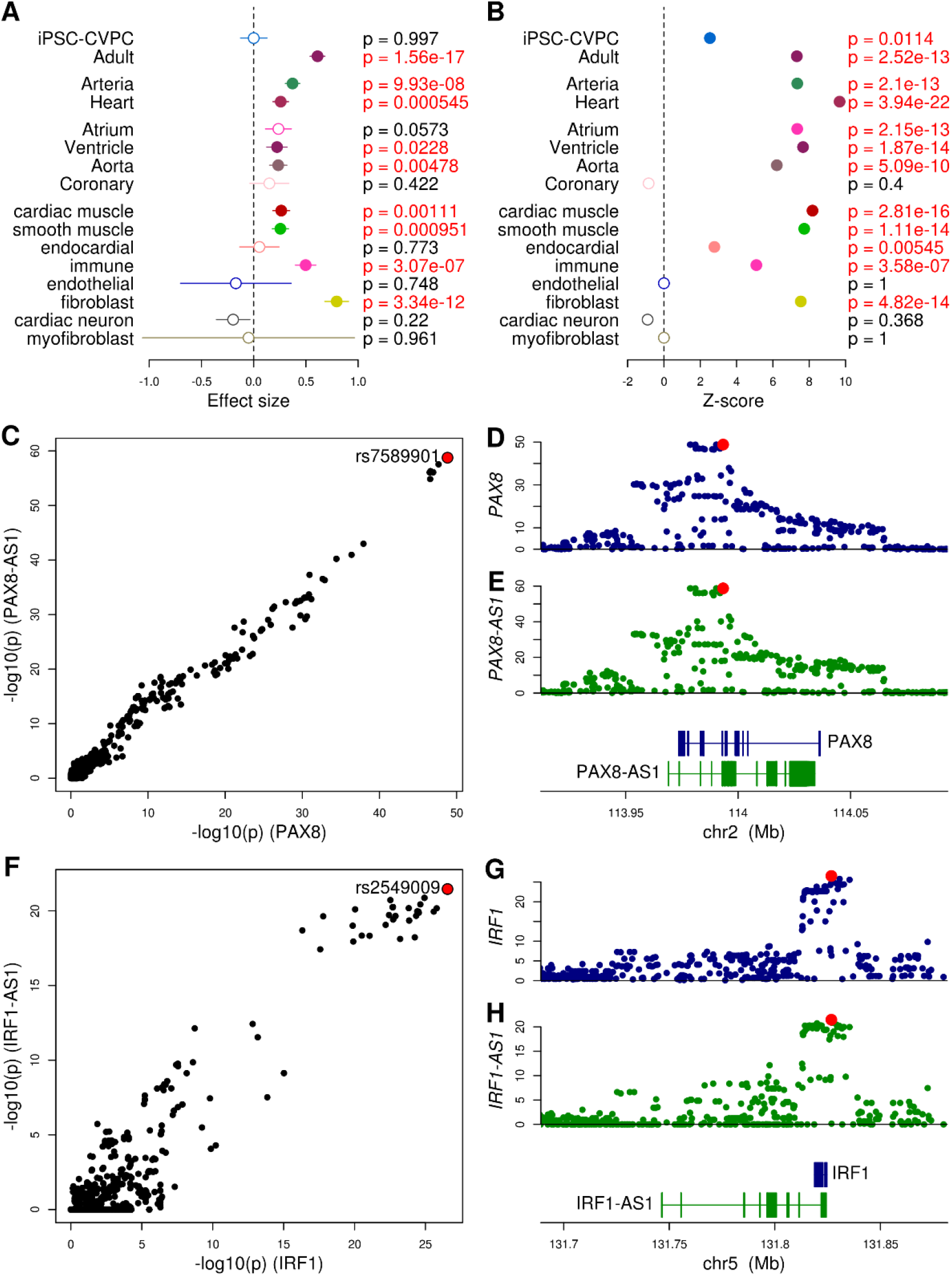
eQTL signals associated with multiple genes. (A) Enrichment of eGenes that share the same eQTL signal with other eGenes or eIsoforms for having stage-, organ-, tissue- and cell type-eQTLs measured by linear regression analysis. Dots represent effect size and segments represent standard errors. (B) eGenes that share the same eQTL signals are enriched for being associated with same stage, organ, tissue or cell type calculated using a permutation test. Dots represent Z-scores. (C-H) eQTL signals shared between a gene and its associated antisense RNA: (C-E) *PAX8* and *PAX8-AS1;* (F-H) *IRF1* and *IRF1-AS1*. (C,F) scatterplots showing the −log_10_ (p-value) for the gene (X axis; A: *PAX8;* D: *IRF1*) and for the antisense RNA (Y axis; A: *PAX8-AS1;* D: *IRF1-AS1*). (D,E,G,H) eQTL signal for the gene (in blue, panels B and E) and for the antisense RNA (in green, panels C and F). The lead eVariant is highlighted in red.

### Cardiac genes and their paired antisense RNA enriched for sharing eQTL signals

Since antisense RNAs are a particular class of non-coding RNAs with regulatory function ^48,49^, we hypothesized that they may share eQTL signals with their associated gene. We observed that 47 pairs of genes and their associated antisense RNA shared the same eQTL signal, which was significantly more than expected (odds ratio = 17.1, p = 2.8 x 10^-37^, Fisher’s exact test). For example, rs7589901, located ~350 bp downstream of *PAX8-AS1* TSS, was the lead eQTL for both *PAX8* and *PAX8-AS1* (Figure 3C-E). *PAX8* plays a pivotal role in cardiomyocyte development, growth and senescence ^50^, while *PAX8-AS1* knockdown impairs cell growth ^51^. We also observed cases where the signal for an eIsoform colocalized with the expression of its associated antisense RNA, such as rs2549009, ENST00000472045.1_2 (*IRF1*) and *IRF1-AS1* (Figure 3F-H). *IRF1* is involved in cardiac remodeling and its overexpression results is associated with cardiac hypertrophy ^52^ and rs2549009 is in LD (R^2^ = 0.764) with rs7734334, a diastolic blood pressure-associated SNP ^53^, suggesting a likely role of the gene pair *IRF1/IRF1-AS1* in driving cardiac remodeling in response to high blood pressure. These results show that cardiac genes and their paired antisense RNAs are enriched for sharing eQTL signals.

### Colocalization identifies potential molecular mechanisms underlying GWAS signals

To examine the extent to which the causal variants underlying cardiac eQTLs are associated with cardiac traits and disease, we performed a colocalization test ^44^ between eQTL signals for eGenes and eIsoforms and the GWAS signals for pulse rate, QRS duration, pulse pressure, atrial fibrillation and myocardial infarction, all obtained from the UK BioBank. For this analysis, we focused on 1,444 eGenes and 919 eIsoforms that overlapped or were in close proximity (<500 kb) with genome wide-significant GWAS SNPs and found that 206 and 125, respectively (331 overall), colocalized with high posterior probability of association (PPA) with at least one GWAS signal (PPA ≥ 0.8, Figure 4, Table S7). Since multiple eGenes may share the same eQTL signal and certain eQTL signals may be associated with both gene expression and isoform usage, we identified 210 independent GWAS signals associated with eQTLs, including 65 that colocalized with multiple eQTL signals (range: 2-9). The vast majority of eQTL-GWAS signal colocalizations were associated with pulse pressure (106 signals) and pulse rate (83), whereas QRS interval, atrial fibrillation and myocardial infraction were associated with fewer than 10 signals each (five, nine and seven, respectively).

**Figure 4:**
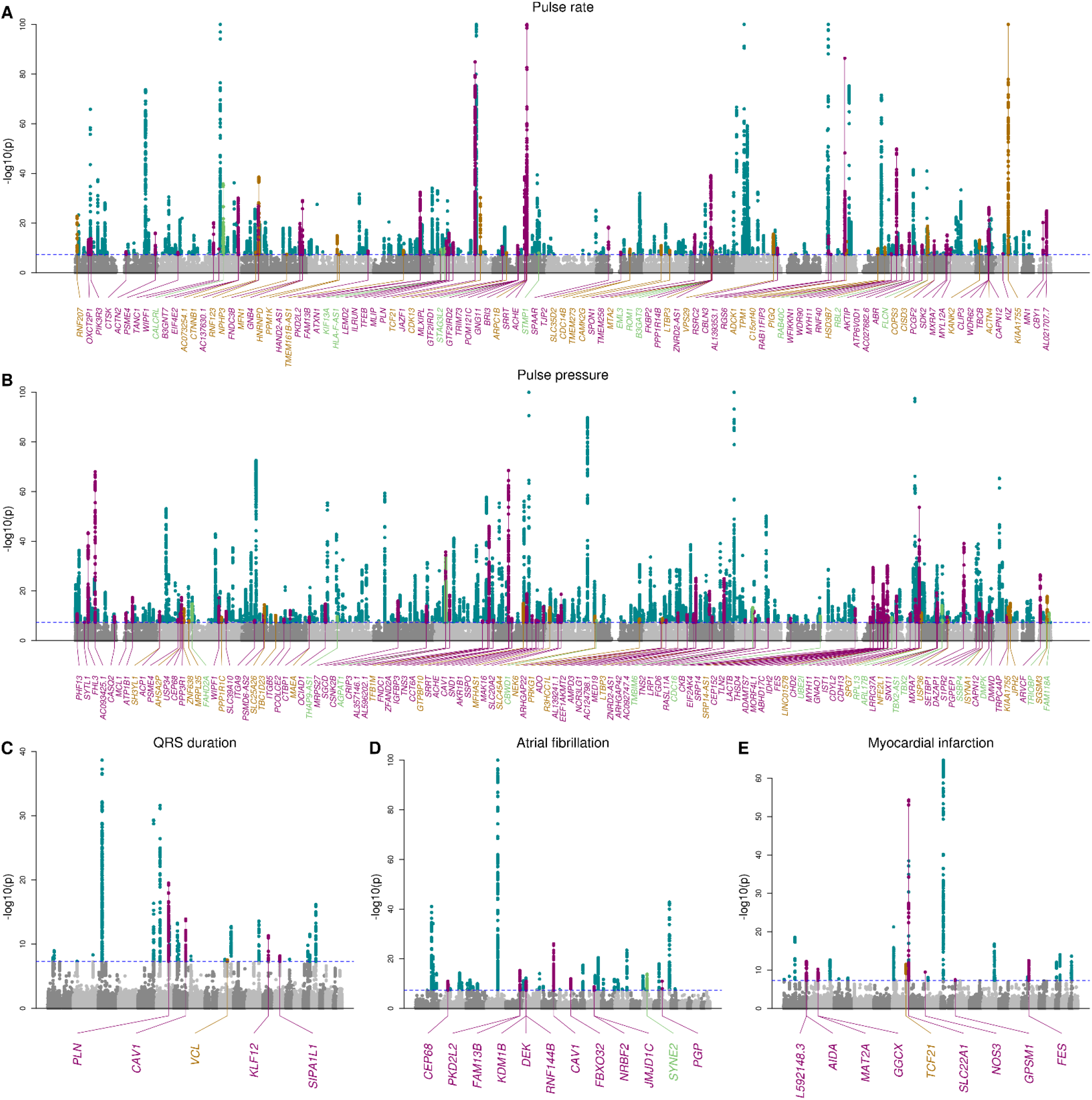
Manhattan plots showing the GWAS signals that colocalize with eQTLs. Manhattan plots showing the GWAS signals for five cardiac traits. Among all genome wide-significant SNPs, the GWAS signals that colocalize with eQTLs for eGenes are highlighted in purple, with eQTLs for eIsoforms (light brown), with eQTL signals for both eGenes and eIsoforms (green), and those that do not colocalize with eQTLs are shown in turquoise. Horizontal dashed blue line represents the genome wide significance threshold (p = 5 × 10^-8^).

Since associations between genetic variation and complex traits can function in a spatiotemporal manner ^24^, we investigated what fraction of the eQTL signals colocalizing with cardiac GWAS signals were associated with developmental stage, organ, tissue or cell type. Of the 206 eGenes and 125 eIsoforms with a high PPA with at least one GWAS signal, we observed that 51 (24.8%) and 21 (16.8%), respectively, had stage, organ, tissue or cell type eQTLs (PPA > 0.8, Figure 5A, Table S7). For example, the cardiac muscle-eQTL signal for *SYNE2* (Figure 2G, Figure 5B-D), a gene that encodes for protein included in the nesprin family that links organelles and nuclear lamina to the actin cytoskeleton ^54^, colocalized with an atrial fibrillation GWAS signal (PPA = 98.3%). Overall, these data show that about one quarter of the eQTL signals that colocalize with cardiac GWAS traits function in a spatiotemporal manner.

**Figure 5:**
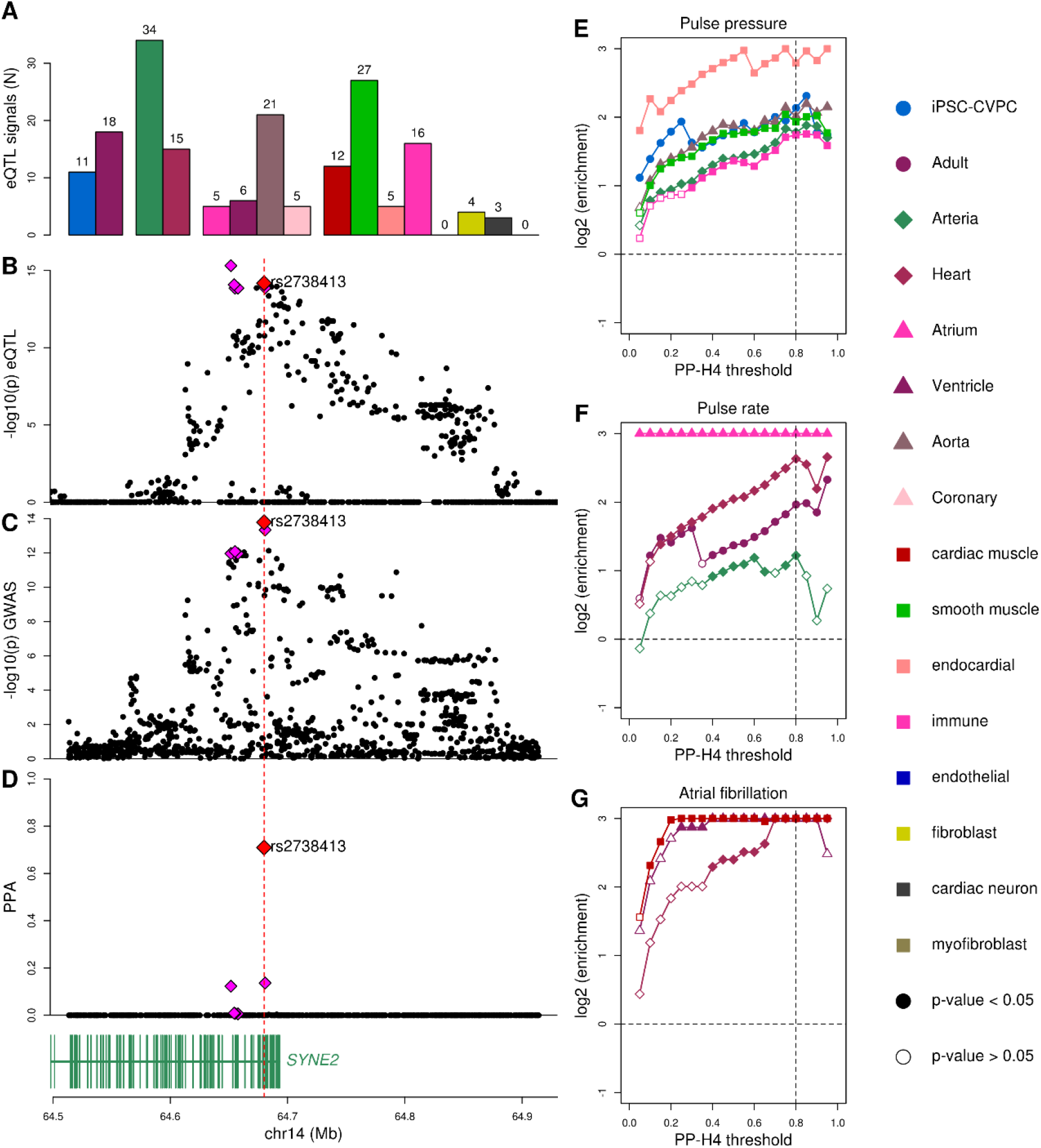
Colocalization between spatiotemporal eQTLs and cardiac GWAS. (A) Barplots showing the number of spatiotemporal eQTL signals that colocalize with cardiac GWAS signals. We observed 51 eGenes and 21 eIsoforms that were associated with one or more spatiotemporal categories colocalize with GWAS traits. Colors represent cardiac stage, organ, tissue and cell type as described in the legend on the right. (B-D) Plots showing (B) cardiac muscle-eQTL signal for *SYNE2*, (C) the GWAS signal for atrial fibrillation and (D) the PPA of each variant in the colocalization. The lead variant (i.e. the variant with highest PPA of being causal for both the eQTL and GWAS signals) is shown as a red diamond: all non-lead variants with PPA > 0.01 are shown as magenta diamonds. (E-G) Line plots showing the enrichment of stage-, organ-, tissue- and cell-type-eQTLs in various GWAS traits: (E) pulse pressure; (F) pulse rate; and (G) atrial fibrillation (ICD10 code: I48). Enrichment is plotted as the log (odds ratio) (Y axis) over all PP-H4 thresholds (0.05 to 0.95, in 0.05 increments) of the eQTL signal colocalizing (0 = not colocalizing; 1 = completely colocalizing) with the GWAS signal (X axis). Only contexts with FDR-corrected p-value < 0.01 at PP-H4 = 0.8 are shown. All associations for the five traits are shown in Figure S5 and Figure S6.

### Cardiac traits enriched for eQTLs that function in specific spatiotemporal contexts

To characterize the associations between each trait and eVariants that function in specific spatiotemporal contexts, we performed an enrichment analysis ^30^. Significant associations were observed for three of the five traits: pulse pressure, pulse rate and atrial fibrillation (Figure S5, Figure S6, Table S8). Pulse pressure was enriched for fetal-like iPSC-CVPC-, arteria-, aorta-, smooth muscle-, endocardial- and immune-eQTLs (Figure 5E); pulse rate was associated with adult -, heart-, arteria-(lower extent), and atrium-eQTLs (Figure 5F); and atrial fibrillation with heart-, left ventricle- and cardiac muscle-eQTLs (Figure 5G). While most of these spatiotemporal enrichments make sense for the given the trait, the enrichment of pulse pressure for fetal-like iPSC-CVPC-eQTLs was surprising (Figure S7). The eQTL signals underlying this enrichment were for five eGenes (*AKR1B1*, *CBWD1*, *RPL13*, *THAP9-AS1* and *TBX2-AS1*) and three eIsoforms (*DMPK*, *RPL13* and *PRKG1*), of which two were iPSC-CVPC-specific (*AKR1B1* and *TBX2-AS1*), indicating that the variants at these loci associated with pulse pressure in the adult could exert their function during cardiac development. Conversely, seven adult-eGenes (*ACHE*, *EIF4E2*, *FLCN*, *RAB40C*, *RBL2*, *SPON1* and *WFIKKN1*) and three adult-eIsoforms (*AC073254.1*, *B3GAT3*, and *RBL2*), of which six were adult-specific (*ACHE*, *EIF4E2*, *RAB40C*, *RBL2*, *SPON1* and *AC073254.1*),colocalized with pulse rate, resulting in the enrichment of pulse rate for adult-eQTLs (Figure S8). The association between pulse pressure and immune cell-eQTLs was also an unexpected observation. Multiple immune cell-eGenes that colocalized with pulse pressure are involved in immune response and inflammation, including *ASAP2*, *SH3YL1*, *ARVCF* and *ATP1B1*^55–58^, and changes in their expression may affect the mechanisms that trigger the inflammatory response and its effects on blood pressure. In summary, by integrating context-associated eQTLs with GWAS, we were able to find that the SNPs associated with three of the five cardiac traits are enriched for being functional in specific spatiotemporal contexts, including associations between pulse pressure and fetal-like iPSC-CVPC-eQTLs and between pulse rate and adult-eQTLs.

### Genetic fine mapping identifies putative causal variants for hundreds of loci

To identify putative causal variants underlying the shared GWAS and eQTL signals we used the colocalization data to conduct genetic fine mapping ^14^. For each of the 210 independent GWAS signals and each of their colocalizing eGenes and eIsoforms (331 GWAS-eQTL combinations) we computed 99% credible sets (defined as the SNPs whose sum of PPAs is >99%). In the 65 cases where multiple eGenes or eIsoforms colocalized with the same GWAS signal, we retained only the credible set with the smallest number of SNPs and, in cases of multiple credible sets with the same number of SNPs, we retained the one having the lead SNP with highest PPA. Across the five cardiac GWAS traits, we found that most credible sets (113, 53.8%) included 10 or fewer SNPs, including 28 (13.3%) having one single causal variant and 51 (24.3%) between two and five (Figure 6A-E, Table S9). Only two credible sets, both associated with QRS duration, included more than 100 SNPs and were considered as not resolved. Fifteen of the 79 credible sets with five or fewer SNPs were associated with spatiotemporal eQTL signals, suggesting that the association between these genetic variants and cardiac traits likely occurs at specific developmental stages, tissues or cell types. Interestingly, fine mapping resulted in credible sets with five or fewer SNPs for three of the pulse pressure GWAS signals that colocalized with fetal-like iPSC-CVPC-eQTLs (for two eGenes: *AGPAT1* and *CBWD1;* and one eIsoform: *RPL13*, Figure 5E, Figure S7). These results show that genetic fine mapping loci containing colocalized GWAS and eQTL signals reduces the number of candidate causal variants to only a handful in the majority of loci and provides a spatiotemporal molecular mechanism underpinning the association between genetic variation and cardiac traits.

**Figure 6:**
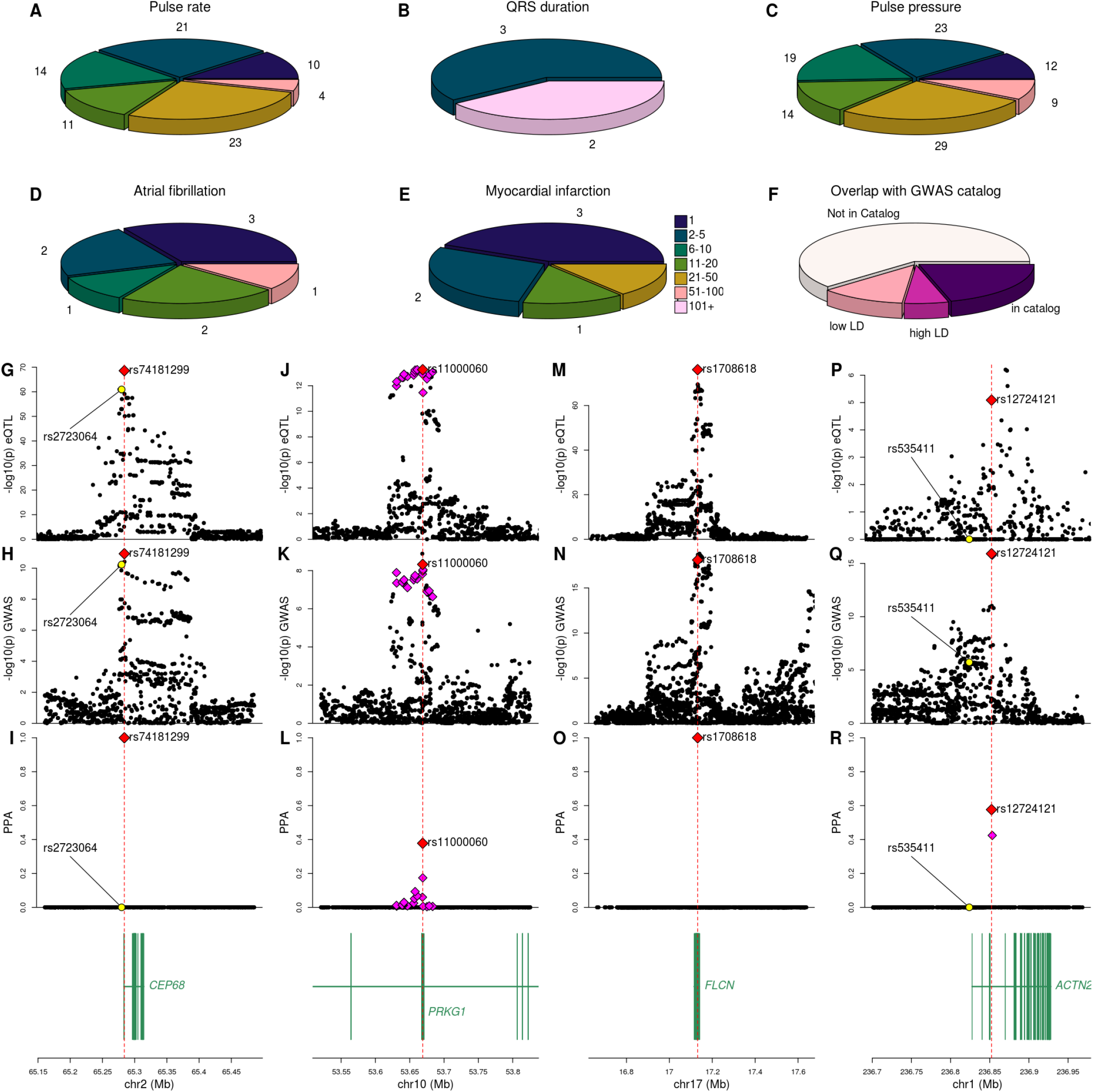
Fine mapping of stage and cell type-eQTLs colocalizing with cardiac traits. (A-E) Pie charts showing for each cardiac trait the distribution of SNPs in 99% credible sets. The colors indicate the number of SNPs and the numbers around the perimeter indicate the number of credible sets. (F) Pie chart showing the overlap between the lead SNP at each GWAS signal in the five cardiac traits and the index SNP for the GWAS signal in the GWAS catalog for the same trait. SNPs in high LD have R^2^ > 0.8 and SNPs in low LD have 0.2 ≤ R^2^ ≤ 0.8. (G-O) Plots showing eQTL signals (top row; panels G, J, M), the GWAS signals (middle row; panels H, K, N) and the PPA of each variant in the colocalization (bottom row; panels I, L, O) at three loci: (G-I) *CEP68* expression and atrial fibrillation; (J-L) *PRKG1* (isoform ENST00000643582.1_1) expression and pulse pressure; (M-O) *FLCN* expression and pulse rate; and (P-R) *ACTN2* expression and pulse rate. The variant with highest PPA of being causal for the colocalized eQTL and GWAS signals is shown as a red diamond. All other variants included in the 99% credible set are shown as magenta diamonds. The location of (G-I) rs2723064 (previously associated with atrial fibrillation in the *CEP68* locus) and (P-R) rs535411 (previously associated with heart failure) are shown in yellow.

To determine if the causal variants identified by our colocalization analysis correspond to lead index GWAS SNPs that have previously been described, we investigated whether they were reported as associated with the same trait in the GWAS Catalog ^1^. We intersected the SNPs with the highest PPA in each of the 210 credible sets with 4,772 trait-associated index SNPs in the GWAS catalog ^1^, and found that only 43 (20.5%) have been previously reported as index SNP for the same trait (Figure 6F, Table S9). For the other 167 signals, we tested their LD with SNPs reported in the GWAS catalog and found that only for 13 signals (6.2%), the lead variant was in high LD (R^2^ > 0.8) and for 25 (11.9%) the lead variant was in low to moderate LD (0.2 ≤ R^2^ ≤ 0.8), indicating that the causal variant is likely not the reported index SNP but rather in LD with it. For the remaining 129 (61.4%), “novel” signals the putative causal SNP that we identified was in a locus not associated with the same trait in the GWAS catalog.

In some cases, our analysis provides a spatiotemporal context for previously identified putative causal variants. For instance, we found that the SNP (rs74181299) with the highest PPA for atrial fibrillation and the cardiac muscle-eQTL for *CEP68* (PPA ≈ 1, Figure 6G-I), is in moderate LD (R^2^ = 0.75) with the previously identified index variant rs2723064 (PPA ≈ 0, 4.2 kb upstream) ^1,59^. We also found that the SNP (rs2738413) with the highest PPA for atrial fibrillation and the cardiac muscle-eQTL in the intron of *SYNE2* (Figure 2G, Figure 5B-D) had been previously identified as an index variant ^2,4^ but proposed to affect expression of the estrogen receptor *ESR2* ^4^, which is located downstream of *SYNE2*, as 17β-estradiol has arrhythmogenic effects on cardiomyocytes ^60^. Since *SYNE2* encodes a nesprin protein, which affects the mechanical properties of the actin cytoskeleton ^61,62^, and *ESR2* is not expressed in cardiac samples, our results suggest that the most likely mechanism underlying the association between rs2738413 and atrial fibrillation involves changes in the expression of *SYNE2* in cardiac muscle cells. Similarly, the SNP (rs11000060) with the highest PPA for pulse pressure and the fetal-like-eQTL for *PRKG1* (isoform ENST00000643582.1_1, PPA = 0.38, Figure 5E, Figure S7, Figure 6J-L) had been previously identified as the index variant ^63^.

On the other hand, many of the fine-mapped GWAS signals were novel (i.e., not in GWAS catalog). For instance, the SNP (rs1708618) with the highest PPA for pulse rate and the eQTL for an *FLCN* isoform (ENST00000389168.6_2) (PPA ≈ 1, Figure 6M-O) had not previously been described in cardiac-related GWAS. However, it has been shown that loss of *FLCN* in the heart results in excess energy and upregulated mitochondrial metabolism, suggesting an important role of this gene in cardiac homeostasis ^64^. The GWAS SNP (rs12724121) with the highest PPA for pulse rate and the eQTL for *ACTN2* (Figure 6P-R) is another example of a novel finding. *ACTN2* encodes alpha-actinin-2, a major component of the sarcomere Z-disc expressed in cardiac and skeletal muscle cells. Rare missense mutations in *ACTN2* result in ventricular fibrillation, cardiomyopathy, and sudden death ^65,66^, and a distal variant (rs535411) associated with heart failure in a GWAS ^67^ was in low R^2^ but high D’ with rs12724121 (D’ = 1; R^2^ = 0.035, 28.5 kb upstream), suggesting a potential synthetic association ^68,69^ exists between these two variants. Our results indicate that rs12724121 may regulate pulse rate (i.e, a proxy for heart rate) through the altered expression of *ACTN2* and, since elevated resting heart rate may cause heart failure ^70^, our findings may explain the previously observed GWAS association between this locus and heart failure. Of note, these observations show the importance of investigating LD in terms of D’, suggesting that synthetic associations ^68,69^ between variants should be considered when characterizing the causal SNPs at GWAS loci. In summary, our fine mapping analysis identified the likely causal SNPs in 81 previously characterized GWAS loci and 129 novel GWAS loci, demonstrating the power of fine mapping approach to uncover insights into the biology of cardiac traits and disease.

## Discussion

We have conducted one of the largest eQTL analyses of human cardiac samples considering both eGenes and eIsoforms and taking spatiotemporal context into consideration. We combined 180 fetal-like cardiac samples (iPSC-CVPC) with 786 adult cardiac samples from multiple tissues, including atrium, ventricle, aorta and coronary artery. We showed that 44.9% of eIsoforms were either not associated with an eGene or had a different eQTL signal than their associated eGene; and that eGenes were enriched for being associated with regulatory variants at promoters and enhancers while eIsoforms were enriched for being associated with variants that affect post-transcriptional modifications. Taking spatiotemporal information about the cardiac samples into account provided us with the unique opportunity to identify more than 2,500 eQTL signals dependent on developmental stage, organ, tissue and/or cell type. Using these data, we show that regulatory variants underlying eQTL signals shared between multiple eGenes are more likely to function in a spatiotemporal manner; and that eGenes which share the same eQTL signals tend to display the same spatiotemporal regulation. Our findings also indicate that fetal-like-eQTLs are less likely to be associated with multiple eGenes than adult-eQTLs, which suggests that gene expression could be more individually regulated in early cardiac development. We demonstrate that, when paired genes and antisense transcripts are both eGenes, they are enriched for sharing the same eQTL signal, suggesting that a tight regulation of protein synthesis is required. Overall, we have generated an invaluable resource comprised of cardiac eQTL signals for 11,692 eGenes and 7,165 eIsoforms, which can be used to understand the molecular mechanisms underlying the association of genetic variants with cardiac traits and disease.

Recent studies showed that certain GWAS loci are associated with tissue- and cell type-specific regulatory elements and eQTLs ^4,15,16,24,30^. Furthermore, GWAS of multiple cardiac traits, such as atrial fibrillation and PR interval, identified loci associated with embryonic development-associated genes and genes whose mutations are known to cause serious heart defects, such as *TTN*, *GATA4*, *MYH6*, *NKX2-5*, *PITX2* and *TBX5*^2–4^ However, the extent to which regulatory variants that function in a spatiotemporal manner underlie cardiac trait GWAS signals was unknown. We showed that many eQTL signals that colocalize with cardiac GWAS traits function in a spatiotemporal manner. Furthermore, we found that three of the five traits examined in this study were enriched for eQTLs that function in specific spatiotemporal contexts. Surprisingly, pulse pressure was enriched for fetal-like-eQTLs, indicating that a subset of genetic variants associated with this adult trait exert their function during early cardiac developmental.

Using colocalization between eQTL and GWAS signals, we fine mapped 210 unique GWAS signals for five cardiac traits and disease. We were able to identify one single likely causal variant (with posterior probability of being causal >99%) for 28 of these GWAS signals, while for additional 51 we were able to restrict the number of likely causal variants to fewer than five. Fifteen of these fine-mapped loci were associated with spatiotemporal eQTLs, including four that were iPSC-CVPC-eQTLs (*AGPAT1*, *CBWD1*, *RPL13* and *PRKG1*). For 81 (38.6%) of the 210 GWAS signals, the SNP with the highest PPA in the credible was either the index SNP (43 signals) or in LD (38 signals) with the index SNP for the same trait in the GWAS catalog. For the remaining 129 (61.4%) GWAS signals, the putative causal SNP that we identified was in a locus not associated with the same trait in the GWAS catalog. These findings show that fine mapping provides the blueprint to both understand the molecular mechanisms underlying known and novel GWAS loci and to uncover insights into the biology of important cardiac traits and disease. Of note, there were hundreds of GWAS signals that did not colocalize with eQTLs, indicating that additional studies will likely require integration with epigenomic data to identify candidate causal variants ^37^. Overall, we show that using our cardiac eQTL resource for fine mapping identified the causal variant underlying hundreds of GWAS signals in five cardiac traits, led to an understanding of the underlying spatiotemporal context, and provided novel insights into the biology of the corresponding cardiac GWAS trait.

## Methods

### Data processing

We obtained RNA-seq and WGS data from two sources: 180 iPSC-CVPC samples from 139 subjects from the iPSCORE Collection ^33,71^ and 786 adult cardiac samples (227 aorta, 125 coronary artery, 196 atrial appendage and 238 left ventricle) from 352 GTEx subjects ^25^.

RNA-seq data for all 966 cardiac samples (180 iPSC-CVPC and 786 adult) was processed as previously described in detail ^29^. Briefly, RNA-seq data from the two sources was obtained from dbGaP (phs000924 and phs000424) and integrated. FASTQ files were aligned to 62,492 autosomal genes and their corresponding 229,835 isoforms included in Gencode V.34lift37 ^72^, as described ^29,33,73^. Only 19,586 autosomal genes with TPM ≥ 1 in at least 10% of samples were considered as expressed and used for eQTL analysis. Likewise, 37,032 isoforms (TPM ≥ 1 and usage >10% in at least 10% of samples) from 10,337 expressed genes were used for isoform eQTL analysis.

We previously described deconvolution of cell type proportions for all 966 cardiac RNA-seq samples in detail ^29^. Briefly, we determined marker genes for eight cardiac cell types (cardiac muscle, cardiac neuron, endocardial, endothelial, fibroblast, immune, myofibroblast and smooth muscle) using single cell RNA-seq ^30,74,75^ and used their average expression levels in each cell type to deconvolute cell type proportions using CIBERSORT ^27^.

VCF files from WGS data were obtained from dbGaP (phs001325 and phs000424). All variants with allele frequency – 5% and in Hardy-Weinberg equilibrium (p > 1 × 10^-6^ in the GTEx samples) were used for eQTL analysis. A kinship matrix was built using plink 1.90b3x ^76^ on variants using the genotype of 1,634,010 SNPs with allele frequency between 30% and 60% in the 1000 Genomes Phase 3 project and genotyped in both GTEx and iPSCORE. This set of variants was also used to generate genotype principal components.

### Covariates for eQTL mapping

To determine the optimal combination of PEER factors to maximize the number of cardiac eQTLs, we selected random 200 genes. To avoid biases due to the expression levels of these genes, we divided all autosomal expressed genes in ten deciles and selected 20 random genes from each decile. We performed eQTL mapping using the following covariates: 1) sex; 2) normalized number of RNA-seq reads; 3) % of reads mapping to autosomes or sex chromosome; 4) % of mitochondrial reads; and 5) 20 genotype principal components. To account for ancestry differences between individuals, we used genotype principal components, rather than reported ancestry. To determine the genotype principal components, we used the same 1,634,010 SNPs with allele frequency between 30% and 60% in the 1000 Genomes Phase 3 project and genotyped in both GTEx and iPSCORE that were used to build the kinship matrix. We merged VCF files from 1000 Genomes, GTEx and iPSCORE and performed PCA using plink 1.90b3x ^77^. We calculated 300 PEER factors ^78^ on the 10,000 expressed genes with the largest variance across all samples and used different combinations (between 10 and 300) to perform the eQTL mapping for each of the 200 test genes. We selected 285 PEER factors to perform gene eQTLs, as this number was associated with the largest number of eQTLs in the 200 test genes. For isoform eQTLs we used 80 PEER factors, as a compromise between obtaining the largest number of eQTLs and computational burden.

### eQTL mapping

For each expressed gene, we used *bcftools query* ^79^ to obtain the genotypes for all the variants within 500 kb of each autosomal gene’s coordinates. To account for relatedness between samples, we performed eQTL mapping using a linear mixed model (LMM) with limix v.3.0.4 ^46^ (*scan* function):

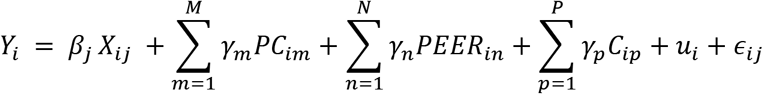

Where *Y_i_* is the normalized expression value for sample *i*, *β_j_* is the effect size (fixed effect) of SNP *j*, *X_ij_* is the genotype of sample *i* at SNP *j*, *PC_im_* is the value of the *m*^th^ genotype principal component for the individual associated with sample *i*, *γ_m_* is the effect size of the *m*^th^ genotype principal component, *M* is the number of principal components used (M = 20), *PEER_in_* is the value of the *n*^th^ PEER factor for sample *i*, *γ_n_* is the effect size of the nth genotype principal component, *N* is the number of PEER factors used, *C_ip_* is the value of the pth covariate for sample *i*, *γ_p_* is the effect size of the pth covariate, *P* is the number of covariates used, *u_i_* is a vector of random effects for the individual associated with sample *i* defined from the kinship matrix, and *∈_ij_* is the error term for individual *i* at SNP *j*. The principal components term captures each individual’s ancestry. The random effects term captures relatedness between samples.

To perform FDR correction, we used a two-step procedure similar to the one described in Huang et al. ^80^. 1) For each gene, p-values were FDR-corrected using eigenMT, which takes into account the LD structure of the tested variants ^81^, and the most significant variant (top hit) was considered as lead-eQTL. If multiple variants had the same p-value, the one with the largest absolute effect size was considered as lead. 2) Across all genes, p-values were corrected for false discovery rate (FDR) using Benjamini-Hochberg’s correction and considered only eQTLs with q-values < 0.05 as significant.

For all significant eQTLs, we tested whether additional variants had independent associations (conditional eQTLs). For each eGene, we regressed out the genotype for the lead eQTL and repeated the eQTL mapping. We repeated this operation to discover up to five conditional associations.

To test the overlap between eQTLs and intergenic regions, introns, promoters, UTRs, splice donor sites, splice acceptor sites and exons, we selected the lead variant for each eGene and eIsoform and tested its overlap with Gencode V.34lift37 genes, promoters (defined as the 2000 bp upstream of the transcription start site), exons and UTRs. For the intronic variants, we calculated their distance from the closest exon, in order to determine their overlap with splice sites.

Summary statistics for all eQTLs are reported in Figshare ^82^.

### Detecting cell type-, stage- and tissue-specific eQTLs

To detect associations between eQTLs and cell types, tissue or developmental stage, we used a linear mixed model with an interaction between genotype and each cell type, tissue or developmental stage using the *iscan* function in limix ^46^:

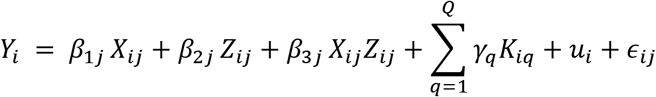

Where *Y_i_* is the normalized expression value for sample *i*, *β*_1*j*_ is the effect size (fixed effect) of SNP *j*, *X_ij_* is the genotype of sample *i* at SNP *j*, *β*_2*j*_ is the effect size of cell type *j*, is the fraction of cell type *j* for sample *i*, *β*_3*j*_ is the effect size of the interaction between genotype *X_ij_* and cell type *Z_ij_*, *K_ip_* is the value of the pth covariate for sample *i*, *γ_q_* is the effect size of the pth covariate, *Q* is the number of covariates used (as defined for the eQTL analysis described above), *u_i_* is a vector of random effects for the individual associated with sample *i* defined from the kinship matrix, and *∈_ij_* is the error term for individual *i* at SNP *j*. We only tested the significant lead QTLs (primary and conditional) for each eGene.

To perform FDR correction, we adjusted p-values of each variant using Bonferroni’s method. We adjusted the p-values independently for each cell type, tissue or developmental stage.

To determine stage, organ and tissue specificity of eQTLs, for each of these elements (for example: left ventricle), we divided samples in two groups (left ventricle and all the other samples) and tested the association between each of these two groups and genotype, using linear regression (*lm* function in R). To calculate cell type associations, for each cell type, we obtained the samples in the top and bottom quartiles and tested the associations between each of these groups and genotype using linear regression. P-values were then corrected using Benjamini-Hochberg’s method.

All possible scenarios based on the significance of the interaction analysis and the two groups are shown in Figure S1 (stage: iPSC-CVPC), Figure S2 (tissue: left ventricle) and Figure S3 (cell type: cardiac muscle).

### Validation of cell type-associated eQTLs

We obtained the relative accessibility score (RAS) of 286,725 snATAC-seq peaks obtained from human adult hearts ^47^. Peak coordinates were lifted over the hg19 genome using liftOver.

To obtain a list of the variants most likely to be causal for each eGene, we performed genetic fine mapping of each eQTL signal using the *finemap.abf* function in the *coloc* R package ^44^. This Bayesian method converts p-values of all the variants tested for a specific gene to posterior probabilities of association (PPA). We selected 201,082 variants with PPA > 0.01, intersected their coordinates with the snATAC-seq peaks using *bedtools intersect*, and found that 18,928 (corresponding to 10,180 eGenes, including 367 associated with cardiac muscle) overlapped snATAC-seq peaks. If multiple variants associated with the same gene overlapped snATAC-seq peaks, we considered only the variant with the highest PPA.

To test if cardiac muscle-associated eQTLs were more likely to overlap snATAC-seq peaks associated with specific cell types, for each of the snATAC-seq cell types (atrial cardiomyocyte, ventricular cardiomyocyte, adipocyte, fibroblast, endothelial, smooth muscle, lymphocyte, macrophage and nervous), we performed a paired t-test between the relative accessibility score (RAS) value for the tested cell type and the mean value across all other cell types for the 367 cardiac muscle-associated eGenes. We repeated this test for all the other cell types. Results for this analysis are shown in Table S5.

### Colocalization between eQTLs signals for eIsoforms and their associated eGene and between different eGenes or eIsoforms

For each of the 5,744 eIsoforms whose associated genes had eQTLs, we performed colocalization using the *coloc.abf* function from the *coloc* package in R ^44^. This Bayesian method uses p-values of the variants tested for two traits (eIsoform and eGene) to calculate the posterior probabilities (PP) of five hypotheses at a specific locus: 1) H0: neither trait has a significant association at the tested locus; 2) H1: only the first trait is associated; 3) H2: only the second trait is associated; 4) H3: both traits are associated but the underlying variants are different; and 5) H4: both traits are associated and share the same underlying variants. Since multiple eQTL signals may be present for both eIsoforms and eGenes (primary and conditional eQTLs), for each eIsoform/eGene pair, we considered only the colocalization with highest PP-H4.

We used the same colocalization approach to determine whether two eGenes or eIsoforms from different genes shared the same eQTL signal.

To determine whether paired genes and antisense RNAs are enriched for sharing an eQTL, we obtained the gene symbols of all eGenes from Gencode. We identified 163 gene/antisense pairs where both the gene and its antisense RNA were eGenes and whose gene symbols were “Gene A” and “Gene-A-AS1”. We found that 43 of these pairs shared the same eQTL signal (PPA ≥ 0.8) and tested whether this proportions was different than expected using Fisher’s exact test.

### Colocalization between GWAS and eQTL signals

We obtained summary statistics from five cardiac traits from the pan-UKBB repository (https://pan.ukbb.broadinstitute.org/). All data was obtained on hg19 coordinates. We sorted and indexed all the files using *tabix* ^79^ and, for each trait, extracted all genome-wide significant SNPs (meta-analysis p-value 5 < 10^-8^). We found 1,444 eGenes and 919 eIsoforms that overlapped or were in close proximity (<500 kb) with genome-wide significant SNPs. We performed colocalization between the eQTL signal of each of these eGenes and eIsoforms and all the genome-wide significant GWAS signals at each locus using the *coloc.abf* function in R ^44^ We considered a GWAS signal as colocalizing with an eGene or eIsoform if PP-H4 > 0.8. All colocalization results between each pair of GWAS and eQTL have been deposited to Figshare ^82^. Since multiple eIsoforms or combinations of eIsoforms and their associated eGene may colocalize with the same trait, underlying the presence of a single gene-trait association, for downstream analyses we considered a trait as “colocalizing” with a gene if PP-H4 was > 0.8 for the eGene or any of its eIsoforms.

To determine the number of unique colocalizations between eQTL and GWAS signals, we computed the LD between each pair of lead SNPs across all 331 colocalizations and obtained clusters of eQTL signals that shared the same lead SNP or that had lead SNPs in high LD (D’ > 0.8). This resulted in 210 unique signals.

### Fine mapping and obtaining 99% credible sets

Using the *coloc.abf* function in R ^44^, we fine mapped the GWAS signals by calculating the posterior probability (PPA) for each tested variant to be causal based on the colocalization between the GWAS and eQTL signals. We next sorted the variants according to their PPA in decreasing order and selected all the variants whose cumulative PPA was 99% as credible sets. For each of the 210 unique signals, we selected the credible set that had the fewest SNPs. If two credible sets had the same number of SNPs, the set containing the lead variant with the highest PPA was chosen.

### Enrichment for the association of cardiac traits with stage-, organ-, tissue- and cell type-eQTLs

Enrichment of the associations was calculated using a Fisher’s Test at multiple PPH4 thresholds (0–0.95; at 0.05 increments), where the contingency table consisted of two classifications: (1) if the variant was significantly context-associated (FDR< 0.05); and (2) if the variant colocalized with the GWAS trait greater than each PP-H4 threshold. We considered as associated all the traits that had FDR-corrected p-value < 0.1 (Benjamini-Hochberg) at the 0.8 threshold.

## Data availability

All data used in this manuscript is available through dbGaP, the Pan UK BioBank resource and Figshare. RNA-seq and genotype information are available for GTEx and iPSCORE through the dbGaP studies phs000424 (GTEx), phs000924 (iPSCORE, RNA-seq) and phs001325 (iPSCORE, whole genome sequencing). RAS values for snATAC-seq peaks were obtained from Supplemental Table 5 in Hocker et al. ^47^. GWAS summary statistics were obtained from the Pan UK BioBank resource (https://pan.ukbb.broadinstitute.org/). The GWAS manifest file was downloaded from https://docs.google.com/spreadsheets/d/1AeeADtT0U1AukliiNyiVzVRdLYPkTbruQSk38DeutU8/edit#gid=511623409. Summary statistics for all eQTLs, fine mapping results and supporting data for all the figures and supplemental tables associated with this manuscript have been deposited at Figshare:https://doi.org/10.6084/m9.figshare.c.5594121.

## Acknowledgements

This work was supported by a California Institute for Regenerative Medicine grant GC1R-06673-B, NSF-CMMI division award 1728497, and NIH grants HG008118, HL107442 and HG011558. JPN and TDA were supported by T15LM011271.

## Author information

KAF, ADC and MD conceived the study. MD, JPN, TDA and HM performed the analyses. HM performed quality check on RNA-seq samples. KAF and ADC oversaw the study. MD and KAF prepared the manuscript.

## References

1. Buniello, A. et al. The NHGRI-EBI GWAS Catalog of published genome-wide association studies, targeted arrays and summary statistics 2019. Nucleic Acids Res 47, D1005–D1012 (2019).

2. Roselli, C. et al. Multi-ethnic genome-wide association study for atrial fibrillation. Nat Genet 50, 1225–1233 (2018).

3. van Setten, J. et al. PR interval genome-wide association meta-analysis identifies 50 loci associated with atrial and atrioventricular electrical activity. Nat Commun 9, 2904 (2018).

4. Nielsen, J.B. et al. Biobank-driven genomic discovery yields new insight into atrial fibrillation biology. Nat Genet 50, 1234–1239 (2018).

5. Loots, G.G. et al. Identification of a coordinate regulator of interleukins 4, 13, and 5 by cross-species sequence comparisons. Science 288, 136–40 (2000).

6. Oh, S. et al. Enhancer release and retargeting activates disease-susceptibility genes. Nature (2021).

7. Edwards, S.L., Beesley, J., French, J.D. & Dunning, A.M. Beyond GWASs: illuminating the dark road from association to function. Am J Hum Genet 93, 779–97 (2013).

8. Cano-Gamez, E. & Trynka, G. From GWAS to Function: Using Functional Genomics to Identify the Mechanisms Underlying Complex Diseases. Front Genet 11, 424 (2020).

9. Ward, L.D. & Kellis, M. Interpreting noncoding genetic variation in complex traits and human disease. Nat Biotechnol 30, 1095–106 (2012).

10. Pennacchio, L.A., Bickmore, W., Dean, A., Nobrega, M.A. & Bejerano, G. Enhancers: five essential questions. Nat Rev Genet 14, 288–95 (2013).

11. Strober, B.J. et al. Dynamic genetic regulation of gene expression during cellular differentiation. Science 364, 1287–1290 (2019).

12. Torres, J.M. et al. A Multi-omic Integrative Scheme Characterizes Tissues of Action at Loci Associated with Type 2 Diabetes. Am J Hum Genet 107, 1011–1028 (2020).

13. Vinuela, A. et al. Genetic variant effects on gene expression in human pancreatic islets and their implications for T2D. Nat Commun 11, 4912 (2020).

14. Mahajan, A. et al. Fine-mapping type 2 diabetes loci to single-variant resolution using high-density imputation and islet-specific epigenome maps. Nat Genet 50, 1505–1513 (2018).

15. Cuomo, A.S.E. et al. Single-cell RNA-sequencing of differentiating iPS cells reveals dynamic genetic effects on gene expression. Nat Commun 11, 810 (2020).

16. Jerber, J. et al. Population-scale single-cell RNA-seq profiling across dopaminergic neuron differentiation. Nat Genet 53, 304–312 (2021).

17. Greenwald, W.W. et al. Subtle changes in chromatin loop contact propensity are associated with differential gene regulation and expression. Nat Commun 10, 1054 (2019).

18. Wilber, A., Nienhuis, A.W. & Persons, D.A. Transcriptional regulation of fetal to adult hemoglobin switching: new therapeutic opportunities. Blood 117, 3945–53 (2011).

19. Viart, V. et al. Transcription factors and miRNAs that regulate fetal to adult CFTR expression change are new targets for cystic fibrosis. Eur Respir J 45, 116–28 (2015).

20. Huang, P. et al. Comparative analysis of three-dimensional chromosomal architecture identifies a novel fetal hemoglobin regulatory element. Genes Dev 31, 1704–1713 (2017).

21. Consortium, E.P. An integrated encyclopedia of DNA elements in the human genome. Nature 489, 57–74 (2012).

22. Roadmap Epigenomics, C. et al. Integrative analysis of 111 reference human epigenomes. Nature 518, 317–30 (2015).

23. Chiou, J. et al. Single-cell chromatin accessibility identifies pancreatic islet cell type- and state-specific regulatory programs of diabetes risk. Nat Genet 53, 455–466 (2021).

24. Kim-Hellmuth, S. et al. Cell type-specific genetic regulation of gene expression across human tissues. Science 369 (2020).

25. Consortium, G.T. The GTEx Consortium atlas of genetic regulatory effects across human tissues. Science 369, 1318–1330 (2020).

26. Bonder, M.J. et al. Identification of rare and common regulatory variants in pluripotent cells using population-scale transcriptomics. Nat Genet 53, 313–321 (2021).

27. Newman, A.M. et al. Determining cell type abundance and expression from bulk tissues with digital cytometry. Nat Biotechnol 37, 773–782 (2019).

28. Wang, X., Park, J., Susztak, K., Zhang, N.R. & Li, M. Bulk tissue cell type deconvolution with multi-subject single-cell expression reference. Nat Commun 10, 380 (2019).

29. D’Antonio, M. et al. In heart failure reactivation of RNA-binding proteins is associated with the expression of 1,523 fetal-specific isoforms. PLoS Comput Biol 18, e1009918 (2022).

30. Donovan, M.K.R., D’Antonio-Chronowska, A., D’Antonio, M. & Frazer, K.A. Cellular deconvolution of GTEx tissues powers discovery of disease and cell-type associated regulatory variants. Nat Commun 11, 955 (2020).

31. Barker, D.J.P. Fetal origins of cardiovascular disease. Ann Med 31, 3–6 (1999).

32. Alexander, B.T., Dasinger, J.H. & Intapad, S. Fetal programming and cardiovascular pathology. Compr Physiol 5, 997–1025 (2015).

33. D’Antonio-Chronowska, A. et al. Association of Human iPSC Gene Signatures and X Chromosome Dosage with Two Distinct Cardiac Differentiation Trajectories. Stem Cell Reports 13, 924–938 (2019).

34. Banovich, N.E. et al. Impact of regulatory variation across human iPSCs and differentiated cells. Genome Res 28, 122–131 (2018).

35. Knowles, D.A. et al. Determining the genetic basis of anthracycline-cardiotoxicity by molecular response QTL mapping in induced cardiomyocytes. Elife 7 (2018).

36. D’Antonio-Chronowska, A., D’Antonio, M. & Frazer, K.A. In vitro Differentiation of Human iPSC-derived Cardiovascular Progenitor Cells (iPSC-CVPCs) Bio-Protocol 10 (2020).

37. Benaglio, P. et al. Allele-specific NKX2-5 binding underlies multiple genetic associations with human electrocardiographic traits. Nat Genet 51, 1506–1517 (2019).

38. Lian, X. et al. Directed cardiomyocyte differentiation from human pluripotent stem cells by modulating Wnt/beta-catenin signaling under fully defined conditions. Nat Protoc 8, 162–75 (2013).

39. Burridge, P.W. et al. Chemically defined generation of human cardiomyocytes. Nat Methods 11, 855–60 (2014).

40. Tohyama, S. et al. Distinct metabolic flow enables large-scale purification of mouse and human pluripotent stem cell-derived cardiomyocytes. Cell Stem Cell 12, 127–37 (2013).

41. Takeda, M. et al. Development of In Vitro Drug-Induced Cardiotoxicity Assay by Using Three-Dimensional Cardiac Tissues Derived from Human Induced Pluripotent Stem Cells. Tissue Eng Part C Methods 24, 56–67 (2018).

42. Consortium, G.T. et al. Genetic effects on gene expression across human tissues. Nature 550, 204–213 (2017).

43. Yang, K.C. et al. Deep RNA sequencing reveals dynamic regulation of myocardial noncoding RNAs in failing human heart and remodeling with mechanical circulatory support. Circulation 129, 1009–21 (2014).

44. Giambartolomei, C. et al. Bayesian test for colocalisation between pairs of genetic association studies using summary statistics. PLoS Genet 10, e1004383 (2014).

45. Charrier-Hisamuddin, L., Laboisse, C.L. & Merlin, D. ADAM-15: a metalloprotease that mediates inflammation. FASEB J 22, 641–53 (2008).

46. Casale, F.P., Rakitsch, B., Lippert, C. & Stegle, O. Efficient set tests for the genetic analysis of correlated traits. Nat Methods 12, 755–8 (2015).

47. Hocker, J.D. et al. Cardiac cell type-specific gene regulatory programs and disease risk association. Sci Adv 7 (2021).

48. Pan, J. & Zhao, L. Long non-coding RNA histone deacetylase 4 antisense RNA 1 (HDAC4-AS1) inhibits HDAC4 expression in human ARPE-19 cells with hypoxic stress. Bioengineered 12, 2228–2237 (2021).

49. Ghafouri-Fard, S., Khoshbakht, T., Taheri, M. & Ghanbari, M. A concise review on the role of BDNF-AS in human disorders. Biomed Pharmacother 142, 112051 (2021).

50. Wu, Y. et al. Pax8 plays a pivotal role in regulation of cardiomyocyte growth and senescence. J Cell Mol Med 20, 644–54 (2016).

51. Huang, C. et al. PAX8-AS1 knockdown facilitates cell growth and inactivates autophagy in osteoblasts via the miR-1252-5p/GNB1 axis in osteoporosis. Exp Mol Med 53, 894–906 (2021).

52. Jiang, D.S. et al. Interferon regulatory factor 1 is required for cardiac remodeling in response to pressure overload. Hypertension 64, 77–86 (2014).

53. Hoffmann, T.J. et al. Genome-wide association analyses using electronic health records identify new loci influencing blood pressure variation. Nat Genet 49, 54–64 (2017).

54. Zhang, X. et al. Syne-1 and Syne-2 play crucial roles in myonuclear anchorage and motor neuron innervation. Development 134, 901–8 (2007).

55. Cao, W. et al. Inducible ATP1B1 Upregulates Antiviral Innate Immune Responses by the Ubiquitination of TRAF3 and TRAF6. J Immunol 206, 2668–2681 (2021).

56. Norton, R.L. et al. Selenoprotein K regulation of palmitoylation and calpain cleavage of ASAP2 is required for efficient FcgammaR-mediated phagocytosis. J Leukoc Biol 101, 439–448 (2017).

57. Yoo, J.Y. et al. LPS-Induced Acute Kidney Injury Is Mediated by Nox4-SH3YL1. Cell Rep 33, 108245 (2020).

58. Abreu-Velez, A.M., Yi, H. & Howard, M.S. Cell junction protein armadillo repeat gene deleted in velo-cardio-facial syndrome is expressed in the skin and colocalizes with autoantibodies of patients affected by a new variant of endemic pemphigus foliaceus in Colombia. Dermatol Pract Concept 7, 3–8 (2017).

59. Christophersen, I.E. et al. Large-scale analyses of common and rare variants identify 12 new loci associated with atrial fibrillation. Nat Genet 49, 946–952 (2017).

60. Yan, S. et al. Bisphenol A and 17beta-estradiol promote arrhythmia in the female heart via alteration of calcium handling. PLoS One 6, e25455 (2011).

61. Balikov, D.A. et al. The nesprin-cytoskeleton interface probed directly on single nuclei is a mechanically rich system. Nucleus 8, 534–547 (2017).

62. Arsenovic, P.T. et al. Nesprin-2G, a Component of the Nuclear LINC Complex, Is Subject to Myosin-Dependent Tension. Biophys J 110, 34–43 (2016).

63. Kichaev, G. et al. Leveraging Polygenic Functional Enrichment to Improve GWAS Power. Am J Hum Genet 104, 65–75 (2019).

64. Hasumi, Y. et al. Folliculin (Flcn) inactivation leads to murine cardiac hypertrophy through mTORC1 deregulation. Hum Mol Genet 23, 5706–19 (2014).

65. Good, J.M. et al. ACTN2 variant associated with a cardiac phenotype suggestive of left-dominant arrhythmogenic cardiomyopathy. HeartRhythm Case Rep 6, 15–19 (2020).

66. Bagnall, R.D., Molloy, L.K., Kalman, J.M. & Semsarian, C. Exome sequencing identifies a mutation in the ACTN2 gene in a family with idiopathic ventricular fibrillation, left ventricular noncompaction, and sudden death. BMC Med Genet 15, 99 (2014).

67. Arvanitis, M. et al. Genome-wide association and multi-omic analyses reveal ACTN2 as a gene linked to heart failure. Nat Commun 11, 1122 (2020).

68. Wang, K. et al. Interpretation of association signals and identification of causal variants from genome-wide association studies. Am J Hum Genet 86, 730–42 (2010).

69. Tam, V. et al. Benefits and limitations of genome-wide association studies. Nat Rev Genet 20, 467–484 (2019).

70. Nanchen, D. et al. Resting heart rate and the risk of heart failure in healthy adults: the Rotterdam Study. Circ Heart Fail 6, 403–10 (2013).

71. Panopoulos, A.D. et al. iPSCORE: A Resource of 222 iPSC Lines Enabling Functional Characterization of Genetic Variation across a Variety of Cell Types. Stem Cell Reports 8, 1086–1100 (2017).

72. Frankish, A. et al. GENCODE reference annotation for the human and mouse genomes. Nucleic Acids Res 47, D766–D773 (2019).

73. DeBoever, C. et al. Large-Scale Profiling Reveals the Influence of Genetic Variation on Gene Expression in Human Induced Pluripotent Stem Cells. Cell Stem Cell 20, 533–546 e7 (2017).

74. Tabula Muris, C. et al. Single-cell transcriptomics of 20 mouse organs creates a Tabula Muris. Nature 562, 367–372 (2018).

75. Butler, A., Hoffman, P., Smibert, P., Papalexi, E. & Satija, R. Integrating single-cell transcriptomic data across different conditions, technologies, and species. Nat Biotechnol 36, 411–420 (2018).

76. Chang, C.C. et al. Second-generation PLINK: rising to the challenge of larger and richer datasets. Gigascience 4, 7 (2015).

77. Purcell, S. et al. PLINK: a tool set for whole-genome association and population-based linkage analyses. Am J Hum Genet 81, 559–75 (2007).

78. Stegle, O., Parts, L., Piipari, M., Winn, J. & Durbin, R. Using probabilistic estimation of expression residuals (PEER) to obtain increased power and interpretability of gene expression analyses. Nat Protoc 7, 500–7 (2012).

79. Li, H. A statistical framework for SNP calling, mutation discovery, association mapping and population genetical parameter estimation from sequencing data. Bioinformatics 27, 2987–93 (2011).

80. Huang, Q.Q., Ritchie, S.C., Brozynska, M. & Inouye, M. Power, false discovery rate and Winner’s Curse in eQTL studies. Nucleic Acids Res 46, e133 (2018).

81. Davis, J.R. et al. An Efficient Multiple-Testing Adjustment for eQTL Studies that Accounts for Linkage Disequilibrium between Variants. Am J Hum Genet 98, 216–24 (2016).

82. D’Antonio, M. Fine mapping spatiotemporal mechanisms of genetic variants underlying cardiac traits and disease. figshare, https://doi.org/10.6084/m9.figshare.c.5594121 (2021).

