## Supplemental information for "Fine mapping spatiotemporal mechanisms of genetic variants underlying cardiac traits and disease"

### Supplementary Figures

#### Figure S1: Examples of associations between eQTL signals and cardiac stage

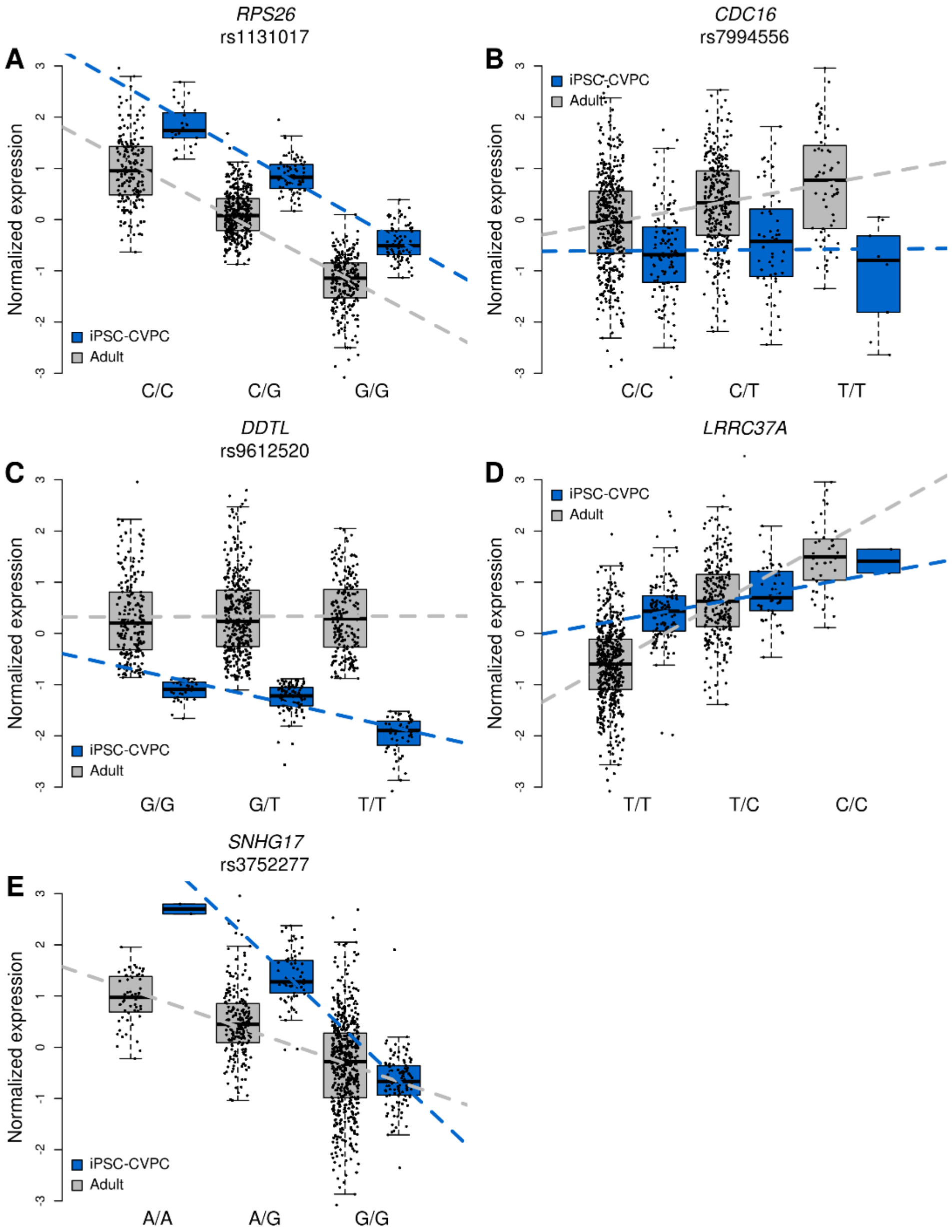


The figure shows examples of five association types between eQTLs and cardiac stage. For each eGene, boxplots describe the normalized expression in iPSC-CVPCs (blue) and all other samples (i.e., adult cardiac samples; gray), grouped by genotype. The panels show examples of: (A) an eGene whose eQTL is shared across both cardiac stages; (B) an eGene whose eQTL is adult-specific: the association between genotype and gene expression is present in the adult cardiac samples but not in iPSC-CVPCs; (C) an iPSC-CVPC-specific eQTL: the association between genotype and gene expression is only present in iPSC-CVPCs; (D) an adult-associated eQTL: although an association between genotype and gene expression is present also in iPSC-CVPCs, it is significantly stronger in the adult samples; and (E) iPSC-CVPC-associated eQTL: while the genotype is associated with gene expression in both iPSC-CVPCs and the adult samples, the eQTL is significantly stronger in iPSC-CVPCs.

#### Figure S2: Examples of associations between eQTL signals and cardiac tissue (adult left ventricle)

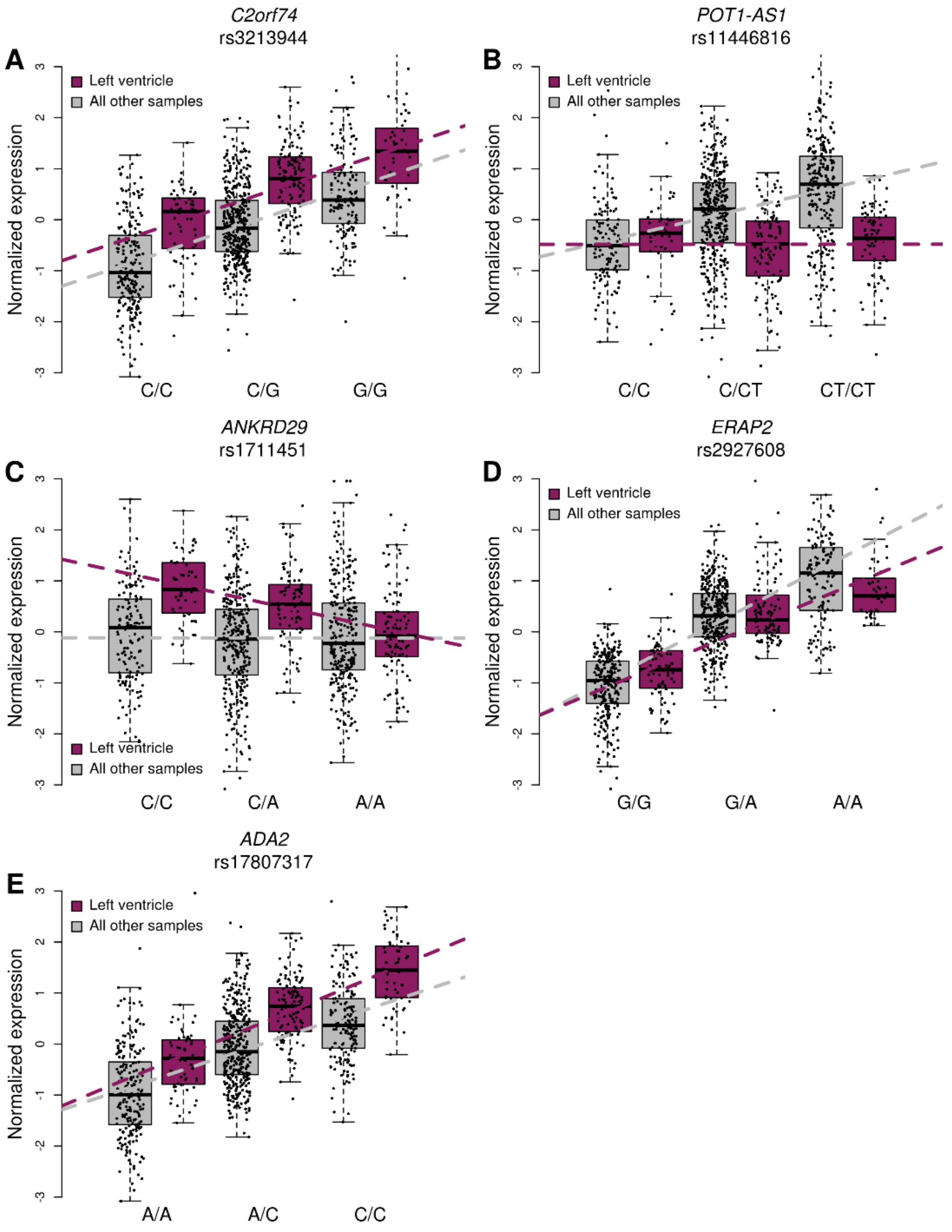


The figure shows examples of five association types between eQTLs and left ventricle. For each eGene, boxplots describe the normalized expression in adult left ventricle (maroon) and all other samples (i.e. all cardiac samples that are not left ventricle; gray), grouped by genotype. The panels show examples of: (A) an eGene whose eQTL is shared across cardiac tissues; (B) an eGene whose eQTL is specific to other cardiac tissues as there is no association between the genotype and gene expression in left ventricle; (C) a left ventricle-specific eQTL: the association between genotype and gene expression is present in left ventricle, but not in the other cardiac tissues; (D) an eQTL that is associated with other cardiac tissues: although an association between genotype and gene expression is present also in left ventricle, it is significantly stronger in the other samples; and (E) left ventricle-associated eQTL: while the genotype is associated with gene expression in both left ventricle and other cardiac tissues, the eQTL is significantly stronger in left ventricle.

**Figure S3: Examples of associations between eQTL signals and cell type (cardiac muscle proportion)**

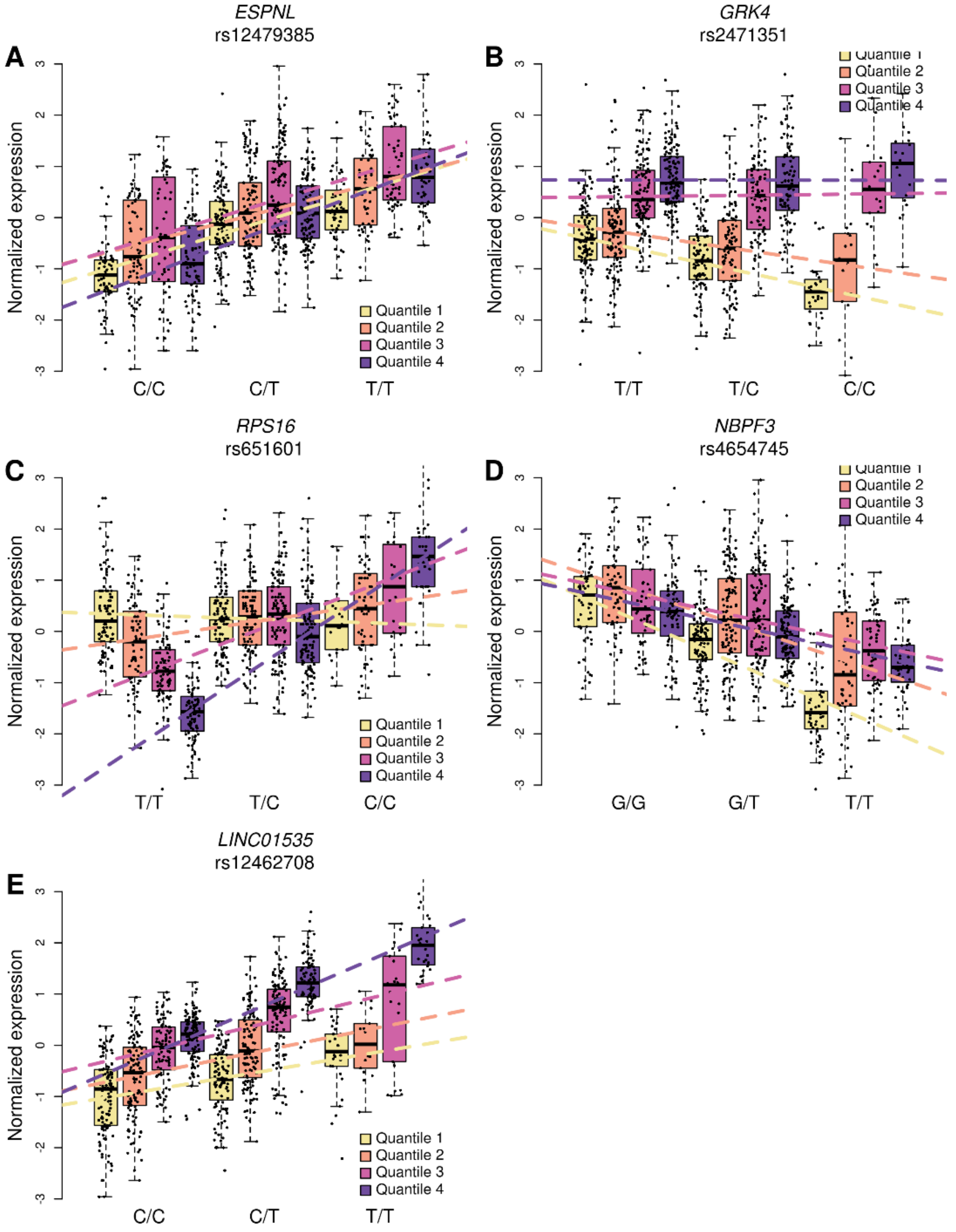


The figure shows examples of five association types between eQTLs and cell types. For each eGene, boxplots describe the normalized expression divided into four quartiles according to their cardiac muscle proportion (yellow = low; purple = high), grouped by genotype. The panels show examples of: (A) an eGene whose eQTL is shared across cell types; (B) an eGene whose eQTL is specific to other cell types but is not associated with cardiac muscle: only samples in the bottom quartiles show an association between the genotype and gene expression; (C) a cardiac muscle-specific eQTL: the association between genotype and gene expression is only present in the top quartiles; (D) an eGene whose eQTL is associated with other cell types: while the association between genotype and gene expression is present in all quartiles, it is significantly stronger in the bottom quartiles; and (E) cardiac muscle-associated eQTL: while the genotype is associated with gene expression in all quartiles, the eQTL is significantly stronger in the top quartiles.

#### Figure S4: Enrichment of cell type- eQTLs for corresponding cell type- snATAC peaks

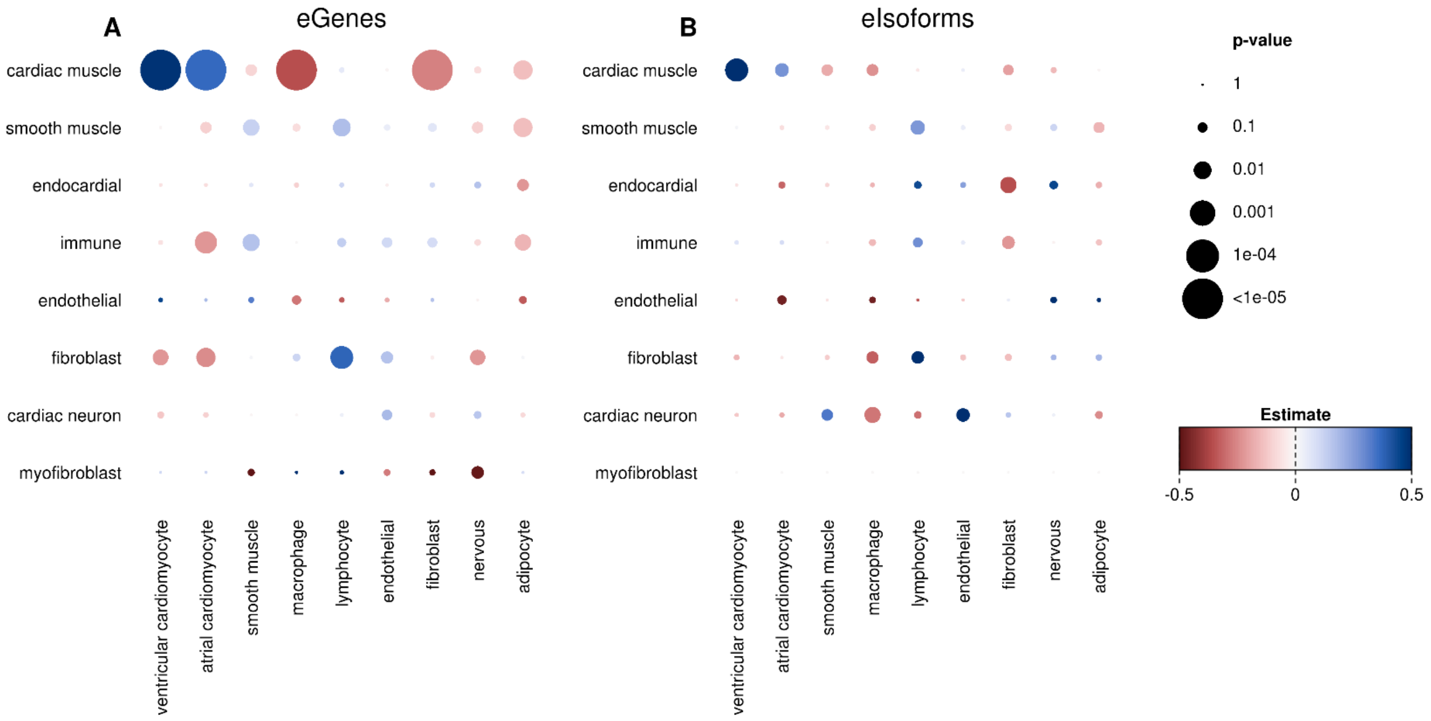


Bubble plots of enrichments of eQTLs associated with each cell type (rows) and the relative accessibility score of nine cardiac cell types obtained from adult cardiac snATAC-seq peaks from an independent study [1]. Estimates and p-values were calculated using the *t.test* function in R.

#### Figure S5: Enrichment of GWAS traits for stage, organ and tissue associations

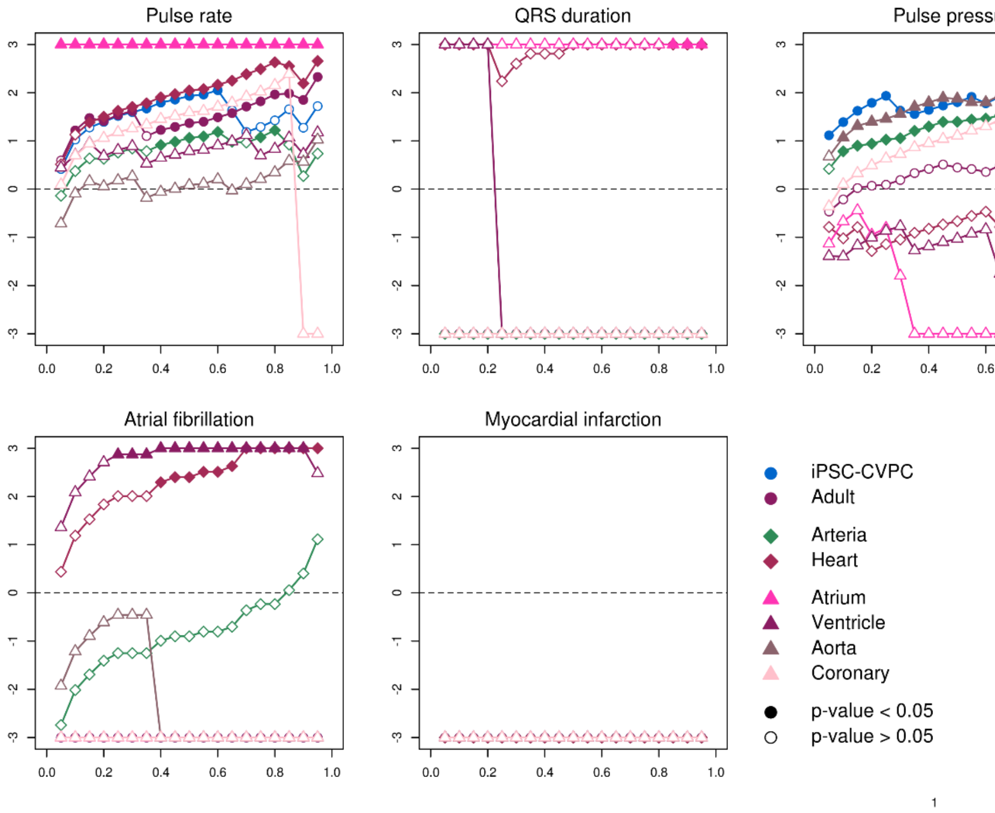


Enrichment for the associations between each of the five cardiac traits and diseases and stage, organ and tissue eQTL associations at multiple PP-H4 thresholds, as described in Donovan et al. [2]. Briefly, at each 0.05 PP-H4 increment between 0.05 and 0.95 (X axis), we tested the enrichment for each stage, organ or tissue for all the eGenes that had PP-H4 above the threshold using Fisher's exact test. The Y axis represents the $\log_2$ of the estimate, as calculated using the *fisher.test* function in R. Filled points represent p-values < 0.05. $\text{Log}_2$ estimate values > 3 or < -3 were set to +3 or -3, respectively. None of the stage, organ or tissue eQTLs colocalize with myocardial infarction signals, therefore all enrichment values are shown on the -3 line.

### Figure S6: Enrichment of GWAS traits for cell type associations

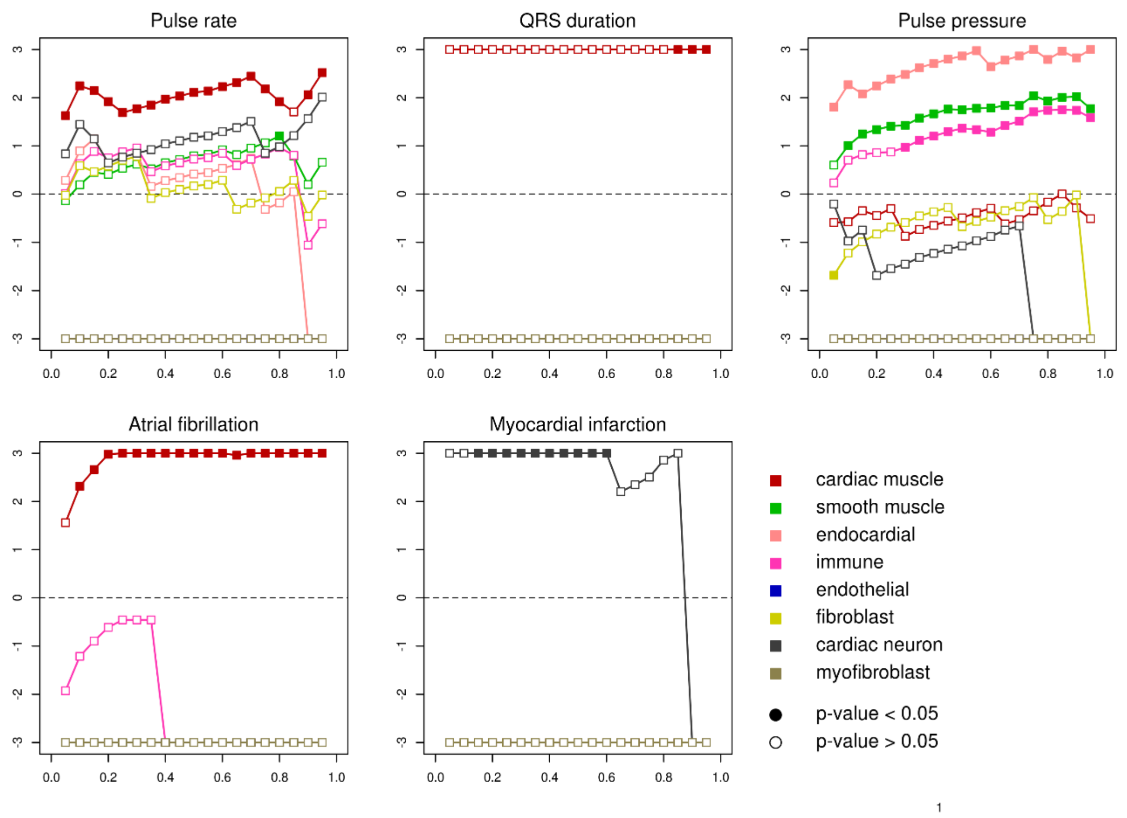


Enrichment for the associations between each of the five cardiac traits and diseases and cell type eQTL associations at multiple PP-H4 thresholds, as described in Donovan et al. [2], as described in Figure S5.

#### Figure S7: Colocalization between iPSC-CVPC- eQTLs and pulse pressure GWAS signals

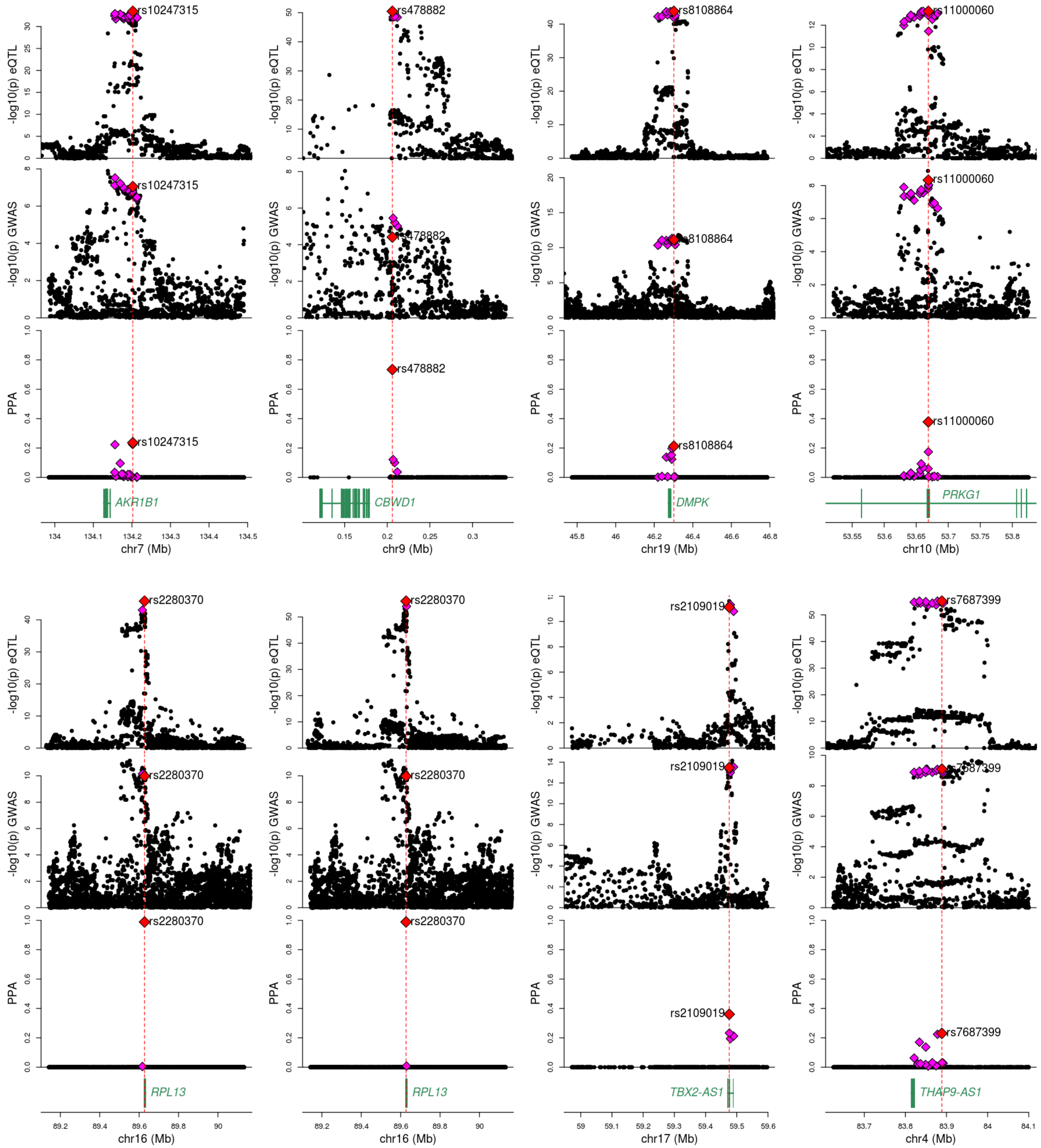


Plots showing iPSC-CVPC- eQTL signals (top row), the colocalizing GWAS signals (pulse pressure, middle row) and the PPA of each variant (bottom row) for five eGenes and three eIsoforms. The iPSC-CVPC-eQTL signal with lead variant rs2280370 is shared between the *RPL13* eGene and one of the *RPL13* eIsoforms (ENST00000565571.5_1). The other two iPSC-CVPC-eQTL signals associated with eIsoforms encode *DMPK* (ENST00000291270.9_3) and *PRKG1* (ENST00000643582.1_1). The lead variant (i.e. the variant with highest PPA of being causal for both the eQTL and GWAS signals) is shown as a red diamond. All non-lead variants included in the 99% credible set are shown as magenta diamonds.

### Figure S8: Colocalization between adult- eQTLs and pulse rate GWAS signals

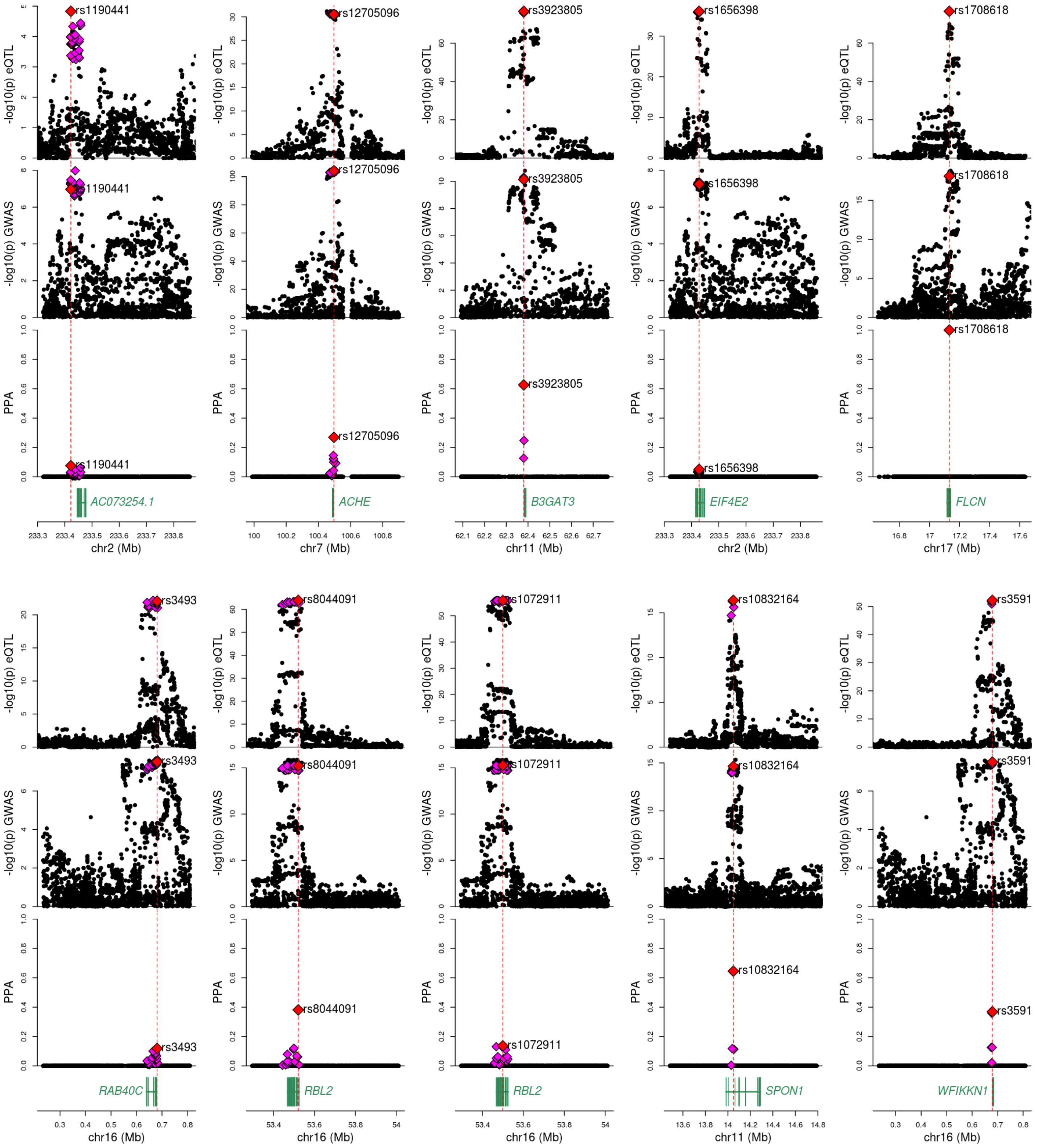


Plots showing adult- eQTL signals (top row), the colocalizing GWAS signals (pulse rate, middle row) and the PPA of each variant (bottom row) for seven eGenes (*ACHE*, *EIF4E2*, *FLCN*, *RAB40C*, *RBL2*, *SPON1* and *WFIKKN1*) and three eIsoforms (*AC073254.1*: ENST00000415506.6_2; *B3GAT3*: ENST00000531383.5_2; and *RBL2*:

ENST00000562850.1_1). The lead variant (i.e. the variant with highest PPA of being causal for both the eQTL and GWAS signals) is shown as a red diamond. All non-lead the variants included in the 99% credible set are shown as magenta diamonds.

#### Supplementary Tables

##### Table S1: Sample metadata

The table shows sample information for all 966 RNA-seq samples which were used for eQTL analysis, including: sample ID (UUID for iPSCORE samples, SRA run ID for GTEx samples), whole genome sequencing ID (UUID for iPSCORE samples, subject ID for GTEx samples); subject ID; subject name; source study (iPSCORE or GTEx); organ (arteria, heart or iPSC-CVPC); tissue (aorta, coronary artery, atrial appendage, left ventricle or iPSC-CVPC); normalized read depth (calculated as the number of reads in each sample divided by the mean number of reads across all samples); % of mitochondrial reads; unique differentiation identifier (UDID, used only for iPSC-CVPC); sex; and cell types deconvoluted using CIBERSORT.

Additional covariates used for the eQTL analysis, including 20 genotype PCs and PEER factors, are reported in Figshare [3].

##### Table S2: eGenes and eIsoforms

The table describes the lead variant for each expressed gene and isoform. For each gene or isoform, shown are: phenotype (gene or isoform), transcript ID (from Gencode V.34lift37; for genes, transcript ID is the same as gene ID), gene ID, gene name, gene type (as defined by Gencode); whether the tested eVariant is primary or conditional, the SNP ID, its chromosome, position, reference allele, alternative allele, SNP ID; beta, standard error of beta and p-value, calculated using limix; number of tests (i.e. number of independent variants tested for the selected gene, calculated using eigenMT) and FDR-corrected p-value calculated by eigenMT; q-value (Benjamini-Hochberg-corrected p-values); and whether the tested gene or isoform is an eGene ("Yes" for all genes and isoforms with q-value ≤ 0.05).

Full summary statistics for all genes and isoforms are reported in Figshare [3].

##### Table S3: Colocalization between each eIsoform and its associated eGene

For each of the 5,744 eIsoforms whose associated gene is an eGene, the table shows all the colocalization between isoform eQTL and gene eQTL signals as calculated using the *coloc.abf* in the R package *coloc*: transcript ID, gene ID, the type of eQTL for the eIsoform and the eGene, the posterior probability of each hypothesis, the variant with the highest posterior probability of being causal and its posterior probability. The five hypotheses are: 1) H0: neither eIsoform nor eGene has a significant association at the tested locus; 2) H1: only the eIsoform is associated; 3) H2: only the eGene is associated; 4) H3: both eIsoform and eGene eQTL signals are associated but the underlying variants are different; and 5) H4: both eIsoform and eGene eQTL signals are associated and share the same underlying variants.

Full PPAs between each pair of eGenes or eIsoforms are reported in Figshare [3].

##### Table S4: Interactions between eQTL signals and stage, organ, tissue or cell type

The table shows all the interactions between eQTLs and stage (iPSC-CVPC or adult), organ (arteria or heart), tissue (atrium, ventricle, aorta or coronary artery) or cell type (cardiac muscle, smooth muscle, endocardial, immune, endothelial, fibroblast, cardiac neuron or myofibroblast). For each eQTL, shown are: phenotype (gene or isoform), transcript ID (from Gencode V.34lift37; for genes, transcript ID is the same as gene ID), gene ID, whether the eVariant is primary or conditional, the SNP ID, the interaction; effect size, standard error, p-value and Bonferroni-adjusted p-value for the interaction between genotype and stage, organ, tissue or cell type; effect size, standard error, p-value and Benjamini-Hochberg-adjusted p-value for the tested stage, organ or tissue, or the top quartile for cell types; effect size, standard error, p-value and Benjamini-Hochberg-adjusted p-value for all the other stages, organs or tissues, or the bottom quartile for cell types; whether the eQTL is associated with stage, organ, tissue or cell type, whether it is specific or associated (as shown in Figure S1, Figure S2 and Figure S3). Only significant interactions are shown. The full table is reported in Figshare [3].

#### Table S5: Enrichment of cell type- eQTLs for cell type-associated snATAC peaks

The table shows the enrichments of eQTLs associated with each cell type and cell type-associated snATAC-seq peaks from an independent study [1]. For each cell type in the eQTL analysis, we performed a paired t-test between the relative accessibility score (RAS) value for each cell type in the snATAC-seq dataset and the mean value across all other cell types for the cell type-associated eGenes. The table shows: phenotype (gene or isoform); cell type in the eQTL analysis and in the snATAC-seq analysis; estimate, 95% confidence interval and p-value calculated using the *t.test* function in R.

#### Table S6: eVariants shared between multiple eGenes or eIsoforms

The table shows the eVariants that are shared between multiple eGenes or eIsoforms. For each eVariant, we indicate the number of associated genes (eGenes or genes with associated eIsoforms) and their gene name.

Full PPAs between each pair of eGenes or eIsoforms are reported in Figshare [3].

#### Table S7: Colocalization between eQTLs and GWAS signals

For each eGene and eIsoform overlapping genome-wide significant GWAS loci, the table shows colocalization PPAs. Specifically, shown are: trait ID, trait name; transcript ID, gene ID, gene name; whether the eVariant is primary or conditional; phenotype ("gene" or "isoform"); the SNP ID (chromosome, position, reference and alternative allele, separated by underscores) and PPA for the SNP with the strongest PPA; and the PPA for each of the five colocalization hypotheses.

#### Table S8: GWAS traits enrichment for stage, organ, tissue and cell type- eQTLs

The table shows the enrichment analysis for the colocalization between stage, organ, tissue and cell type- eQTLs and each of the five GWAS traits: trait ID, trait name (as in Table S7); tested interaction, as described in Table S4; estimate, confidence interval, $\log_2$ ratio of the estimate and p-value, as calculated using the *fisher.test* function in R. the data described in this table was used as input to create Figure 5E-G, Figure S5 and Figure S6.

#### Table S9: Fine mapped cardiac GWAS loci

The table shows each of the 331 colocalizations between eQTL and GWAS signals. Since multiple eQTL signals may map to the same GWAS signal, we calculated LD between each pair of lead SNPs and obtained 210 clusters of eQTL signals (all SNPs in each cluster had D' > 0.8). For each of these clusters, we selected the eQTL signal associated with the smallest 99% credible set and used it to fine map its corresponding GWAS signal. This table reports all these colocalizations and the information about 99% credible sets associated with each colocalization: trait ID, trait name; phenotype ("gene" or "isoform"); transcript ID, gene ID, gene name; whether the tested variant is primary or conditional; the PPA for each of the five colocalization hypotheses; the SNP ID, RS ID and PPA for the SNP with the strongest PPA; the number of variants in the 99% credible set; the LD cluster ID (chromosome and cluster, separated by underscore); whether the colocalization was used for fine mapping; RS ID of the SNP in highest LD in the GWAS catalog (frozen at June 2, 2021) [4], its $R^2$ and D' values, calculated using LDlink (https://ldlink.nci.nih.gov/), the Pubmed IDs associated with the SNP in the GWAS catalog and the category of association with the GWAS catalog. These categories are (Figure 6F): 1) "in catalog", if the SNP with the highest PPA has previously been found in other GWAS for the same trait; 2) "high $R^2$", if the SNP with the highest PPA is in high LD ($R^2 \geq 0.8$) with a SNP associated with the same trait in the GWAS catalog; 3) "high D'", if the SNP with the highest PPA is in high LD ($D' \geq 0.8$ and $R^2 < 0.8$) with a SNP associated with the same trait in the GWAS catalog; and 4) empty, if the SNP with the highest PPA is not in high LD with any SNP associated with the same trait in the GWAS catalog.

Full information about each colocalization, including the summary statistics of the GWAS at each locus, the associated eQTL summary statistics, the PPA of each tested SNP and the composition of each credible set, has been deposited to Figshare [3].

### References

1. Hocker, J.D. *et al.* Cardiac cell type-specific gene regulatory programs and disease risk association. *Sci Adv* **7**(2021).
2. Donovan, M.K.R., D'Antonio-Chronowska, A., D'Antonio, M. & Frazer, K.A. Cellular deconvolution of GTEx tissues powers discovery of disease and cell-type associated regulatory variants. *Nat Commun* **11**, 955 (2020).
3. D'Antonio, M. Fine mapping spatiotemporal mechanisms of genetic variants underlying cardiac traits and disease. *figshare*, https://doi.org/10.6084/m9.figshare.c.5594121 (2021).
4. Buniello, A. *et al.* The NHGRI-EBI GWAS Catalog of published genome-wide association studies, targeted arrays and summary statistics 2019. *Nucleic Acids Res* **47**, D1005-D1012 (2019).
